# Dated public evidence and known genes for retrospective evaluation of gene prioritization

**DOI:** 10.64898/2026.11.04.764446

**Authors:** Chao-Hui Huang, Yujia Wang

## Abstract

Retrospective evaluation of gene prioritization needs evidence that was public before the evaluated genes became known. Public resources are open and reproducible, but are distributed as current releases that also contain evidence produced after the genes became known. This Data Descriptor presents per-gene evidence tables for four areas (oncology, inflammatory bowel disease, metabolism and psoriasis) from public resources. Every feature carries a time axis, the date of its release in a calendar of 183 archived releases of 20 sources, and a leakage axis, a provenance class (A to E) with two risk grades that define five feature sets. Nineteen lists of known genes are dated by the first trial, study, release or classification that supports each gene. They cover targets of a kind of drug, targets of drugs tested in a disease, and genes associated with a disease. The oncology lists also carry the first day the trial registry showed the trial.

## 1 Background & Summary

Methods that rank genes are asked different questions. Which genes will become targets of a kind of drug, such as an antibody–drug conjugate? Which genes will become targets of drugs tested in a disease? Which genes are associated with a disease? Such methods are often trained on known genes and tested on genes held out at random. Public resources such as Open Targets [1], STRING [2], the Human Protein Atlas [3], DepMap [4] and integrated knowledge graphs [5, 6] are a common source of the inputs. They are open and cover many kinds of evidence, and anyone can repeat an analysis that uses them. Method development and benchmarks that others must be able to check therefore rest on them. They do not replace the data that a company generates for its own programmes. Such data are designed for a question, in that the assay, the material and the comparison are chosen for it. They can include results that were never published, and the company controls their use. Public resources hold what has been published or deposited, and how much they hold for a gene depends on how much the gene has been studied [7, 8]. A value that follows the amount of study can raise the apparent performance of a method without carrying biological information. This was shown for the multifunctionality of genes in guilt-by-association analysis [9].

A separate problem arises when public resources are used to test a method retrospectively. They are distributed as current releases, and a current release contains evidence that exists because a gene became known. Drug–target relations and clinical-trial records are created by drug development, GWAS entries by the study of a disease, and the literature on a gene grows once the gene is studied. The timing of this evidence has been measured in the Open Targets Platform. For the 433 novel drug targets approved since 2000, clinical-trial evidence clusters around the year of first approval, and the peaks of most other evidence types moved from after approval to before it around 2015 [10]. A preprint that fitted a classifier of drug approval on 26,278 target–disease pairs reported that the literature-mining features alone carried most of its performance, which its author read as consistent with temporal leakage from publications that follow approval [11]. A model tested on a held-out gene can therefore use evidence that was produced after that gene became known. This follows from how the releases are distributed and says nothing about the quality of the sources. Leakage between training and test data is known to inflate the performance of knowledge-graph models in biomedical link prediction [12]. Evaluations that fix the inputs at a date and judge a method on what is reported afterwards exist for gene prioritization and for protein function. One compared eight prioritization tools on 42 recently reported disease–gene associations before the underlying databases were updated, and found that cross-validation on retrospective data was likely to be overoptimistic, because some data sources are contaminated with knowledge of the associations [13]. In the first CAFA experiment, predictions of protein function were submitted before a deadline and judged on the annotations that accumulated during the following 11 months [14]. Each such evaluation fixes one date. The dataset described here keeps the inputs of many dates, so that a method can be replayed at any quarter end from 2016 to 2024. Studies of the genetic support of drug targets began with the archive of genetic evidence as it stood at the time of analysis [15]. Later studies asked whether support that existed before a programme predicts its success. Support did so prospectively for Mendelian traits [16], and its effect was largely unaffected by the year in which the genetic evidence was discovered [17]. Such questions need inputs that are restricted to a date, which public releases do not supply by themselves.

A historical replay fixes a cutoff date and restricts every input to what was public before it. A model is trained on the genes known at the cutoff and tested on genes that reached a defined milestone later, such as a first trial, an approval or a first reported association. Assembling such inputs by hand is error-prone, for four reasons. Sources date their releases in different ways (a release note, a folder or file timestamp, a publication date). The timestamps of a download server can reflect a later re-upload and not the release. Integrated knowledge graphs inherit dates only through their upstream sources. Some sources keep no earlier release at all.

The dataset (Fig. 1) annotates every evidence component along two axes. The time axis is the release date of the sources from which the component is built (Table 5). For a cutoff date, a snapshot uses the latest release of each source on or before that date, and a component whose sources have no dated release is left out. The leakage axis is a provenance class and two risk grades (Table 3). The class runs from measured biology, which would hold had the gene never been a target, to therapeutic-history evidence, which exists only because the gene was targeted. The two grades record how strongly a value follows the volume of study of a gene and how strongly it encodes therapeutic history. Five feature sets are defined by the grades, from the full set to a biology-only set that contains measured biology alone. A feature set and a cutoff together define a table of features and the dated labels of the known genes. Technical Validation reports the number of features and genes available for each combination.

**Figure 1:**
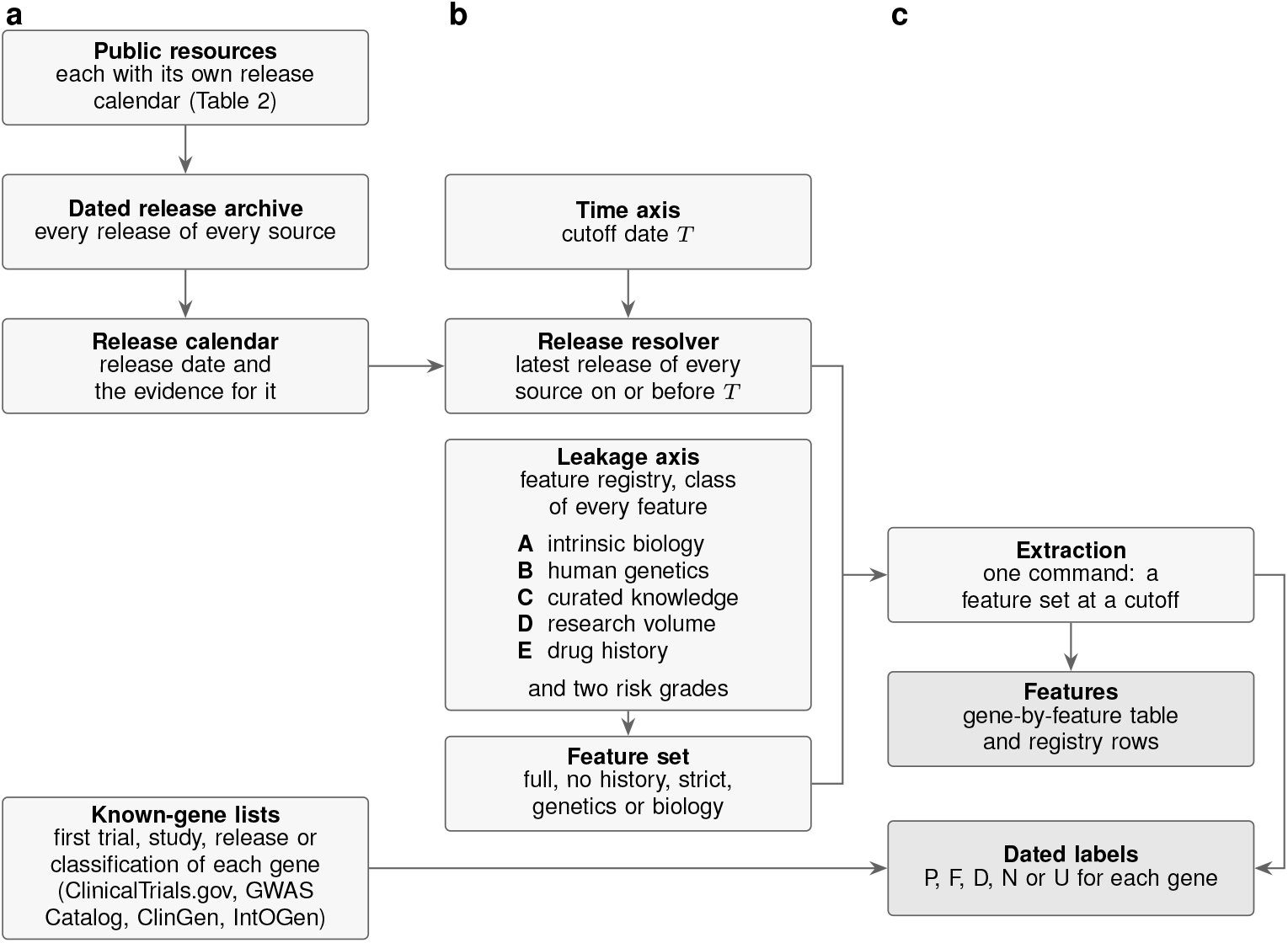
Construction of the dataset and extraction of features and dated labels. **a**, Every public resource is down-loaded as a dated release archive. The release calendar records the release date of every release and the evidence for that date. The lists of known genes record the first trial, approval, study, release or classification that supports each gene and, for the oncology lists, the date on which the trial registry first showed the trial. **b**, The time axis takes a cutoff date, and the release resolver selects the latest release of every source on or before it. The leakage axis gives every feature a provenance class (A, intrinsic biology; B, human genetic evidence; C, curated knowledge; D, research volume; E, drug history) and two risk grades in the feature registry. A feature set (full, no history, strict, genetics or biology) admits the features whose grades are within its caps. **c**, One extraction command applies both axes and returns the selected features and a label for every gene. The labels are P (present: known on or before the cutoff), F (future: known only after it), D (undated gene), N (curated negative) and U (unlabeled). The tables, the manifest and the licence table form the deposit.

The deposit contains the label, feature and gene-set tables of each of four areas, the lists of known genes with the dates from which each gene counts (the first trial, study, release or classification that supports it), the papers and trials behind these genes with their dates, and the curation audits of the lists that rest on trials. It also contains extractions at fixed cutoffs, the feature dictionary with its classification rules, and the release calendar of the source archive. The calendar gives the date of each archived release and the evidence for it. A manifest records the licence of every file. The upstream release archives are not deposited, and the accompanying code downloads and rebuilds them.

Several public datasets support machine-learning work on drugs, their targets and the genes of diseases. Integrated biomedical knowledge graphs combine public resources into one graph for drug repurposing and precision medicine [5, 6, 18, 19]. Each graph is distributed as a snapshot of its sources at the time it was built. Curated drug– target resources record compound–target pairs with the clinical stage of each compound. ChEMBL [20] is the largest open example, and one dataset provides code that regenerates its compound–target pairs from a given ChEMBL release [21]. Other resources hold experimentally validated drug–target links [22] and mechanistic paths from a drug to the disease it treats [23]. Clinical-trial records have been assembled into datasets for predicting trial outcomes [24], and a literature knowledge graph links dated publications, patents and clinical trials [25]. Resources of gene-level features serve ranking methods directly. Pharos and its database TCRD aggregate information on targets and were built for understudied proteins [26]. The Harmonizome holds about 72 million associations between genes or proteins and their attributes, processed from over 70 resources [27]. DrugnomeAI combines 324 gene-level features from 15 sources to estimate the druggability of every protein-coding gene, with models by disease type and by modality [28]. OpenBioLink is a benchmark for link prediction in biomedical networks [29], and its software can also split edges by time. In the present dataset each feature carries the date of the release of its source, and each known gene the date from which it counts. Individual sources such as Open Targets and DepMap keep their earlier releases, each on its own calendar [1, 4]. The Open Targets Platform has also timestamped single pieces of evidence by the date of the primary source or of the curation, which allows the evidence for a target–disease association to be reconstructed for past years [10].

Archived releases of single sources are the starting point of the dataset. They do not supply four things that a retrospective evaluation needs. The first is a release date for every source, with the evidence for it and a rule for sources that have none. The second is one cutoff that applies to all sources together. The third is a provenance class and risk grades, which allow features that record research or drug development to be removed without removing measured biology. The fourth is a list of known genes with the date from which each gene counts. The dataset described here provides these four things. It aligns the releases of its sources by date and assigns each feature a class and two grades. It dates each known gene by the first evidence for it: for a gene targeted by a drug, its first qualifying clinical trial (and, in oncology, the date the trial registry showed the trial), and for a gene–disease association, the study, release or classification that supports it. Each list has a claim, which is either *target* (a drug acts on the gene) or *association* (the gene is linked to a disease), and a scope, which is a modality or a disease.

The data allow four kinds of work. A gene-prioritization method can be benchmarked retrospectively at a chosen cutoff, for drug targets of a modality or an indication and for genes associated with a disease. The same method can be compared across cutoffs or across feature sets. The provenance classes allow an ablation in which one class of features is removed. The release calendar and the dated lists of known genes can be used in other temporal evaluations. A new disease area is added with an area definition file, and the user supplies its lists of known genes and any curation it needs.

## 2 Methods

### 2.1 Input data

Table 2 lists the source resources, the releases used, the basis on which each release is dated and whether the processed records are redistributed. Every archivable source is downloaded into a dated release archive `(raw/<sor>/<release>/)`. The date of each release is stored with its evidence. The evidence is a release note or changelog entry, a folder or file timestamp, a publication date, or a version name that encodes a calendar quarter or year. Release counts, access dates and file sizes are in the release calendar (Data Records). The Open Targets Platform [1] is the default knowledge-graph source. The other integrated graphs [5, 30, 6] are optional and are selected at rebuild time.

**Table 1:** Extraction of one list of each area at the cutoff 2020-02-03. Features: number of features of the full set; A/B/C/D/E: their number in each provenance class. P and F: genes known on or before the cutoff and genes known after it.

| Area | List | Features | A/B/C/D/E | P | F |
|---|---|---|---|---|---|
| Oncology | ADC.trial | 52 | 31/1/17/2/1 | 49 | 16 |
| IBD | IBD.trial | 53 | 31/1/18/2/1 | 111 | 25 |
| Metabolism | T2D.gwas-rep | 53 | 31/1/18/2/1 | 110 | 498 |
| Psoriasis | PSO.gwas-rep | 53 | 31/1/18/2/1 | 43 | 59 |

**Table 2:** Source resources. Releases: number, and month of the first and last release (the exact dates and the evidence for them are in the release calendar). Deposit: partly (the Li et al. 2021 features and the KEGG gene sets are left out), dates only (the AACT copies give the registry dates but are not features), list only (the list of known genes made from the source is deposited, and the source files are not), no (an optional graph rebuilt by the user). Provenance class of the features: class A for HPA, GTEx, DepMap, gnomAD, the Open Targets RNA-expression score, CPTAC, the half-life and internalization tables, the Cell Surface Protein Atlas, UniProt, AlphaFold DB, TED, TCGA and CELLxGENE Census; class B for the Open Targets genetic-association score; class C for MSigDB, the network aggregates, the Open Targets curated-relation evidence (pathways, animal models, somatic mutations) and the glycosylation and disulfide-site counts; class D for the Open Targets literature and connection-count evidence; class E for the Open Targets known-drug evidence, the Tahoe-100M drug counts and the drug-count aggregates (Table 6). Sources with no dated release (TED, TCGA, CELLxGENE Census, Tahoe-100M and PrimeKG) are available only when requested at a cutoff.

| Source | Evidence used | Releases | Dating basis | Licence | Deposit |
|---|---|---|---|---|---|
| Human Protein Atlas [3] | RNA expression, localization | 10; 2017-08 – 2025-11 | file timestamp | CC BY 4.0 | yes |
| HPA blood proteome [33] | plasma proteins | 7; 2020-07 – 2026-06 | file timestamp | CC BY 4.0 | yes |
| GTEx [32] | normal-tissue RNA expression | 6; 2015-10 – 2026-01 | release table | GTEx Portal license | yes |
| STRING [2] | interaction channels | 5; 2015-04 – 2026-08 | file timestamp | CC BY 4.0 | yes |
| DepMap [4] | CRISPR dependency, expression | 25; 2018-08 – 2026-09 | repository record | CC BY 4.0 | yes |
| gnomAD [45, 46] | gene constraint | 4; 2016-08 – 2024-04 | publication date | CC0 | yes |
| CPTAC [47] | protein abundance (oncology) | 1; 2023-05 | file timestamp | CC BY 4.0 (PDC) | yes |
| Protein half-lives [48, 49, 41] | protein turnover in cells | 3; 2018-02 – 2021-10 | publication date | CC BY 4.0; Elsevier | partly |
| Cell Surface Protein Atlas [50] | surface proteins | 1; 2015-04 | publication date | CC0 | yes |
| Internalization [51] | surface-protein endocytosis | 1; 2020-06 | publication date | CC BY 4.0 | yes |
| AACT [34] | registry copies (trial dating) | 39; 2017-03 – 2026-09 | snapshot date | public domain | dates only |
| ClinGen [37] | gene–disease validity (known-gene list) | 1; file created 2026-10-16 | classification date | CC0 | list only |
| IntOGen [36] | cancer driver genes (known-gene list) | 5; 2014-12 – 2024-09 | release name | CC0 | list only |
| MSigDB [52] | gene sets | 15; 2017-04 – 2026-01 | file timestamp | CC BY 4.0; KEGG restricted | partly |
| GWAS Catalog [35] | associations (annotation table; GWAS lists of known genes) | 3; 2019-12 – 2026-09 | release directory date | EMBL-EBI terms | yes |
| UniProt [53] | annotations | 41; 2016-03 – 2026-06 | release notes | CC BY 4.0 | yes |
| Open Targets [1] | evidence by datatype; drug indications and mechanisms | 39; 2016-04 – 2026-09 | folder timestamp | CC0 | yes |
| Monarch [30] | gene–phenotype | 14; 2023-09 – 2026-09 | folder name | CC0; sources keep their terms | no |
| Hetionet [5] | integrated graph | 1; 2017-02 | file timestamps | per node and edge | no |
| DRKG [54] | integrated graph | 1; 2020-06 | file timestamps | Apache-2.0 | no |
| PrimeKG [6] | integrated graph | fetched 2026-09-27 | none | CC0 | no |
| AlphaFold DB [31] | predicted structures | 6; 2021-07 – 2025-09 | changelog | CC BY 4.0 | yes |
| TED [55] | structural domains | fetched 2026-09-28 | none | CC BY 4.0 | yes |
| TCGA PanCanAtlas [56], retrieved through cBioPortal [57, 58] | tumour cohorts (oncology) | cohort tables | none | open access (GDC); cBioPortal: ODbL 1.0, with attribution | per-gene statistics |
| CELLxGENE Census [59] | single-cell specificity (oncology) | Census 2025-11-08 | none | CC BY 4.0 | yes |
| Tahoe-100M [60] | drug metadata (oncology) | revision in source.json | none | CC0 (dataset card) | yes |

Five sources have no dated release and are used only when a user requests them at a cutoff: PrimeKG, TED, TCGA, CELLxGENE Census and Tahoe-100M. Five further sources are archived with their release dates but are not features. They are the SURFY predicted surfaceome (licence CC BY-NC-ND, kept for rebuilding and not redistributed), the Pharos target development levels, DGIdb drug–gene interactions, PubTator3 gene mentions and iCite article years. The licence of each is recorded in the supplementary release calendar. All sources are public.

The lists of known genes are made from two further sources (Table 2): the ClinGen gene–disease validity table and the IntOGen compendia, besides the GWAS Catalog, AACT and Open Targets. They are downloaded into `raw/lists/src/`, and each gene is dated by the records of its source (Lists of known genes and labels). They are not part of the release calendar of the feature sources, and the deposit holds the lists made from them and not the source files.

The GTEx data were obtained from the GTEx Portal on 10/06/2026 (releases V6, V6p, V7, V8, V10 and V11). The deposited tables use the most current release (V11). Tables that a user rebuilds for an earlier cutoff use earlier releases and do not reflect the most current data of the portal, as the GTEx Portal data licence requires us to state. Single-cell tables of ten further cancers can be added from TISCH2, which states no licence. The deposited tables are built without them, from the CELLxGENE Census tables of eight cancers alone.

### 2.2 Feature construction

The build turns each source into columns of per-gene tables indexed by gene symbol. There is one feature table per area (`gene_leaves_*.parquet`) and one metadata row per feature. Expression from the Human Protein Atlas and GTEx enters as summaries across tissues: the maximum nTPM or TPM, the number of tissues above a threshold and the breadth of expression. DepMap enters as gene-level summaries of dependency and expression across cell lines. Open Targets enters as association scores between 0 and 1 for each data type, aggregated over diseases. MSigDB enters as the number of gene sets of each collection that contain the gene. UniProt and AlphaFold enter as summaries of topology, localization and structure (counts, lengths, fractions and confidence). gnomAD enters as constraint metrics. The neighbour and pathway-partner features average one of these features over the neighbours of the gene in a graph or a pathway. A value that a source does not supply stays empty, and the tables are not imputed. The feature dictionary describes every feature in words (`documentation/feature_dictionary.tsv`), and the build code (`build_layer/`) specifies fields, units and transformations. Genes are keyed by the symbol of the gene universe of the area, and the crosswalk maps symbols to accessions (Identifier harmonization).

### 2.3 Evidence components and provenance classes

No source is classified as a whole. Each source is split into evidence components, defined by source, sub-source, entity type and relation or attribute type. A single question assigns each component to one of five provenance classes (Table 3): would the fact still exist if nobody had ever tried to drug the gene? The question was written for the lists of drug targets. It keeps its use for the lists of the claim association, because evidence of drug development and of research volume can enter the features of any list. The evidence that dates an association list needs one more rule. The genetic evidence behind a GWAS list is itself class B, and the features that are built from the evidence of a list are named for each kind of list (`same_source_features`; Lists of known genes and labels). Class A is measured biology, such as omics, structure and population constraint. Class B is human genetic evidence, variants linked to traits (GWAS, ClinVar, gene-burden tests), which was measured in people and does not depend on drugs or trials. Class C is curated biological knowledge: it is biological, but curators have organized it. Class D is research-volume evidence, whose size follows how much a gene has been studied (literature and connection counts; see Background). Class E is therapeutic history, which exists only because a drug was developed or tested.

**Table 3:** Provenance classes assigned to evidence components. The class and the two risk grades determine the feature sets of Table 4.

| Class | Definition | Examples |
|---|---|---|
| A | Measured biology; would hold had the gene never been developed as a target or studied in a disease | RNA expression, CRISPR dependency, sequence, structure, localization, population genetic constraint |
| B | Human genetic evidence; variants linked to traits, measured in people, independent of drugs and trials | Open Targets genetic association (GWAS, ClinVar, gene-burden evidence) |
| C | Curated biological knowledge; biological, but organized by curators | GO terms, gene sets, pathways, phenotypes, curated relations, aggregates over protein-interaction networks, site annotations |
| D | Research-volume evidence; its size depends on how much a gene has been studied | literature co-occurrence, text-mined relations, publication and connection counts |
| E | Therapeutic history; exists only because a drug was developed or tested | drug–target relations, approvals, clinical trials, mechanism of action, drug counts, drug-perturbation signatures |

These are conceptual categories. The class of an implemented feature is the one the rule table assigns (Table 6); for example, an MSigDB gene-set count is class C, because curators choose which genes a set holds.

Each component also receives two risk grades. The study-risk grade measures how strongly the value of the component follows the number of studies on the gene. The history-risk grade measures how strongly the value reflects past drug development. Each grade is none, low, medium or high. A rule table assigns class and grades by source and, for knowledge-graph features, by edge type (`documentation/classification_rules.tsv`). The reason for the assignment of each feature is in `documentation/feature_dictionary.tsv`. The author curated the rule table, and it is applied to every feature of a source; no feature is annotated individually. Table 6 shows how representative components map to classes and grades. Class A features are measured genome-wide by a platform, so their values do not depend on whether a gene was pursued as a target, and they receive low study risk. The exceptions are features whose detection depends on abundance, which the grade does not capture. The protein-interaction network (STRING) and the GWAS Catalog are not features. The network enters only through neighbour and pathway-partner aggregates, which are graded at least class C with medium study risk, and the GWAS Catalog enters only through a display table and the genetic lists of known genes (Lists of known genes and labels). A feature of an unrecognized source receives class C with medium grades. This default keeps the feature out of the biology-only set, which is conservative by design.

### 2.4 Dating of evidence

A feature is admitted at a cutoff only if its public availability before that date can be shown. The evidence is the release calendar. A feature becomes available at the latest of the first dated releases of the sources from which it is built. A snapshot at a cutoff uses, for every source, the latest release published on or before the cutoff (database-level replay). Records within a release are not dated individually; they share the date of the release. Missing dates are not imputed. A source without a dated release gives its features no date, and these features are excluded unless a user requests them. Integrated knowledge graphs are dated through their own releases [5, 6], and a graph without a dated release is treated as undated.

### 2.5 Temporal confidence

The feature registry gives each feature a temporal source, a date basis and a temporal confidence (Table 5). A feature built from dated releases has the temporal source `historical_snapshot`. A feature from a source without a dated release has the temporal source none and the confidence `unavailable`. The confidence follows the kind of evidence that dates the first release of each source of the feature. This evidence is the date basis, read from the date evidence recorded with the release. The weakest kind among the sources of a feature decides its confidence. In the feature dictionary of 144 features, 71 have high confidence, 29 medium, 14 low and 30 are unavailable.

### 2.6 Release resolver

For a cutoff, the resolver selects for each source the latest release published on or before that date. It uses a release calendar compiled from the release notes, changelogs, archives and file timestamps of the sources, with the date evidence recorded (Data Records). Sources that did not exist at the cutoff, such as predicted-structure databases before their first release [31], are omitted from that cutoff. When a source reports only a version number, the release date comes from the strongest available evidence: a release note or changelog entry, the folder or file timestamps on the download server, or the publication date. The calendar records the evidence used, and no date is imputed without it.

Some download servers show timestamps of a later re-upload and not of the release. In that case the release is dated by the source’s own release record. The GTEx releases [32] are dated by the data-release table of the GTEx Portal, the analysis-methods document of the release and its announcements, because the storage timestamps of the files date from a 2023 migration. The opposite case also occurs. A file that a source re-uploads after its release is dated by its own timestamp when that is later than the release. The blood-proteome table of the Human Protein Atlas [33], available from release v19, is dated by its file, so it is never admitted before it can be shown to have been public.

### 2.7 Feature sets

Five feature sets select components by their risk grades (Table 4). The *full* set admits every component. The *no history* set caps the history grade and so drops every component whose value encodes past drug development. The *strict* set caps both grades and so also drops research-volume evidence. The *biology* set admits only components with low study risk and no history risk. In the present rule table these are exactly the class A components, which are measured omics, structure and constraint. The *genetics* set adds the class B components, human genetic evidence, which has medium study risk and no history risk. Curated gene sets and the protein-interaction network contain therapeutic edges, so the biology and genetics sets restrict the derived views accordingly. The selection depends only on the risk grades and not on the cutoff. A feature set is defined at every cutoff, although the features it holds depend on which sources existed at that date (Technical Validation).

**Table 4:** Feature sets along the leakage axis. Each set caps the two risk grades (none = 0, low = 1, medium = 2, high = 3); a component is admitted when both of its grades are at or below the caps, and every set is available at every cutoff.

| **Feature set** (profile name) | history_risk **cap** | study_risk **cap** | **Curated gene sets** | **Effect** |
|---|---|---|---|---|
| Full (full) | high | high | admitted | every component; all derived views |
| No history (no_history) | low | high | excluded | drops components that encode past therapeutic development |
| Strict (strict) | low | medium | excluded | additionally drops research-derived evidence that tracks study volume |
| Biology (biology) | none | low | excluded | measured biology only (class A); derived views that mix therapeutic edges in are dropped |
| Genetics (genetics) | none | medium | excluded | classes A and B: measured biology and human genetic evidence; the same derived views are dropped |

**Table 5:** Temporal confidence of a feature in the feature registry: the kind of evidence that dates the first release of its sources (`date_basis`), the weakest kind counting. A feature with a source that has no dated release has temporal source none, confidence `unavailable`, and is excluded unless explicitly requested.

| Confidence | Date basis | Evidence that dates the release | Examples; direction of a possible error |
|---|---|---|---|
| high | release_record | the provider’s own release notes, changelog, release table or repository record | UniProt, GTEx, DepMap, AlphaFold DB; dated by the provider |
| medium | file_timestamp | the server dates of the files or folders | HPA, MSigDB, STRING, CPTAC, Open Targets folders; a re-upload makes the date late, and a file can also predate public access, so the date is a proxy for it |
| medium | publication | the publication date of the paper that released the data | half-life, surface-protein and internalization tables; a proxy for the date the data were public |
| low | version_name | the month or quarter encoded in the version name, the end of the period used | early Open Targets releases; the date is late by construction |

**Table 6:** How representative components map to provenance classes and risk grades (study risk, history risk). The rule applied is the source rule or, for knowledge-graph features, the edge-type rule of `documentation/classification_rules.tsv`.

| Component (example feature) | Class | Study | History | Reason recorded |
|---|---|---|---|---|
| DepMap CRISPR dependency (depmap__pancan_dependency) | A | low | none | measured omics |
| HPA tissue expression (hpa__n_tissues_ge10) | A | low | none | measured expression |
| Protein topology and predicted structure (struct__log_length) | A | low | none | sequence topology, predicted structure |
| Glycosylation and disulfide sites (struct__n_glycosylation) | C | medium | none | site annotation, partly literature-curated |
| MSigDB gene-set counts (msigdb__n_c6_all) | C | medium | low | curated gene-set membership |
| Open Targets genetic association (opentargets__genetic_association) | B | medium | none | human genetic evidence: variants linked to traits |
| Network neighbours (nbr__depmap__pancan_dependency) | C | medium | none | aggregate over curated, study-dependent knowledge |
| Open Targets literature and connection counts (opentargets__literature, __n_disease) | D | high | medium, low | text-mined literature; connection counts grow with study |
| Open Targets known drug (opentargets__known_drug) | E | high | high | drug–target record |
| Tahoe-100M drugs targeting the gene (tahoe__n_drugs_targeting) | E | high | high | exists because compounds were tested |

### 2.8 Extraction by feature set and cutoff

The extraction applies both annotations. The user chooses a feature set (Table 4), may restrict it to some provenance classes (class A alone, or classes A and B), and may set a cutoff date. Without a cutoff, the features come from the current releases. With a cutoff, a data root is assembled from the releases that the resolver selects, the per-gene features are rebuilt from these releases, and a feature is admitted under the rule of Dating of evidence. Protein-topology features are computed from the UniProt release in use at the cutoff. Predicted-structure features are admitted only from the first AlphaFold DB release onward. The data root for a cutoff is built once and reused by later extractions that resolve to the same releases.

For a list of known genes, the extraction labels each gene by the date from which it counts (`known_from`, defined in Lists of known genes and labels). The label P (present) marks a gene that is known on or before the cutoff and is used as a training positive. The label F (future) marks a gene that becomes known after the cutoff and is held out. The label D (undated) marks a gene without a dated record; it is held out of training unless the user requests otherwise. A gene all of whose trials were withdrawn (registered, but never enrolling a patient) can be left out of the positives on request and is then treated as an ordinary gene. Curated negatives are labelled N, and every other gene is labelled U (unlabeled). The output has four parts: a gene-by-feature table, the label table, the registry rows of the selected features (class, risk grades and availability date), and a JSON record of the settings and of the release resolved for each source.

### 2.9 Lists of known genes and labels

The deposit holds 19 lists of known genes in four areas (Table 8). A list is defined by two choices. Its claim is *target* when a drug acting on the gene was tested, and *association* when the gene is linked to a disease or trait and no drug is involved. Its scope is a kind of drug (a modality) or a disease. The combinations answer three questions about a gene. Which genes become targets of a kind of drug (the ADC, TCE and IO lists)? Which genes become targets of drugs tested in a disease (the inflammatory bowel disease lists)? Which genes come to be associated with a disease or trait (the GWAS, ClinGen and cancer driver lists)? All lists are built from public sources only, and each gene carries the date from which it counts as known.

The lists belong to four areas. An area fixes the gene universe (the 20,151 protein-coding genes of the Human Protein Atlas in all four), the lists that are shown with it, an eligibility annotation, and the curated negatives (oncology only). Oncology is organized by modality (ADC, TCE and IO) and has 32 cancer types as diseases. The inflammatory bowel disease, metabolism (type 2 diabetes, obesity and lipid traits) and psoriasis areas are organized by disease. The positive label of the oncology area is the union of its trial lists (`column pos_ANY_PUBLIC` of the label table), and that of each other area the union of its preset lists (`pos_IBD, pos_T2D, pos_PSO`). The lists differ in biology and in eligibility: ADC targets are cell-surface proteins, and TCE and IO targets need not be, and a GWAS list of a disease follows the genetics of that disease. Evaluation per list is therefore recommended, with a union as a pooled secondary analysis.

#### Lists of the claim target

The trial lists of oncology are chosen by modality. A candidate gene of the antibody–drug-conjugate (ADC) and T-cell-engager (TCE) lists must have an extracellular region in UniProt (GPI-anchored proteins are included, HLA genes are excluded). For each modality, a candidate becomes a positive in one of three ways. Two or more interventional ClinicalTrials.gov studies name it in an intervention of that modality. Exactly one such study names it, and a manual audit of the trial record accepts it. Or a drug of that modality that acts on the gene has a clinical trial, and a public drug record documents the link between drug and gene. These records are the NCI Drug Dictionary, ADC Review and publications (`curation/oncology_drug_evidence.tsv`). Eighteen ADC targets are in the list in this third way, because no trial record names them as a target. Their source field reads curated drug.

The immuno-oncology (IO) list is chosen by mechanism and not by a drug type that a text search can name. A gene is in the list when a clinical-stage drug that modulates it so that the immune system attacks a tumour has an interventional cancer study. Such drugs are checkpoint inhibitors, costimulatory agonists and innate checkpoint blockers, among other classes. The drugs come from a hand-reviewed table shipped with the dataset (`curation/io_targets.tsv`). Each gene is dated by the first such study.

#### Dates of the trial lists

Each gene of a trial list carries the date of its first qualifying trial (`first trial`) and the ClinicalTrials.gov identifier of that trial (`first nct`). A withdrawn trial never enrolled a patient, so it is passed over when a later trial exists. A target all of whose trials were withdrawn is marked `withdrawn only`. The dates of the 18 targets come from ClinicalTrials.gov records, except that of CD5. CD5 is a ricin immunotoxin from before trial registration, and it is dated by the 1991 publication of its phase I trial in cutaneous T-cell lymphoma. Audits corrected three dates (CD5, EPCAM and TPBG). The automatic dates of these three came from an unrelated vaccine trial and from two antibody fusion proteins that are not ADCs. Single-trial hits that no audit reviewed enter the lists (1 of the 65 ADC targets and 9 of the 31 TCE targets), and their `audit` field reads “single trial, not reviewed”. The manual audits ship with the dataset (`curation/`, two tables with 97 records in all). A registry shows a trial as it is today. A trial registered after it started, or a record edited later to name the target, would date a target before anyone could have seen its trial. Each gene of an oncology trial list therefore also carries a registry date (`registry_date`, with `registry_nct`). This is the earliest date on which ClinicalTrials.gov showed one of the qualifying trials of the gene. It is found by applying the matching rules to each of 39 quarterly copies of the registry (AACT [34], 2017-03-09 to 2026-09-01), as that copy recorded each study. A trial counts from its first-posted date when it was a new registration, and otherwise from the date of the first copy whose record qualifies. A trial first posted before the first archived copy (2017-03-09) counts from its first-posted date, which the registry records for every trial. Every oncology gene with a registered trial therefore has a registry date.

The label table records the later of `first_trial` and `registry_date as known_from`. A target counts as known only when its trial has started and the registry shows it. This is a conservative milestone and not the earliest public disclosure of the target. It is the default of the extraction (`--target-dating registry`), because a target should not be a training positive before anyone could have seen its trial. The option `--target-dating trial start` dates targets by `first_trial` alone. The names of the options that contain the word target (`--target-dating, --include-undated-targets, --exclude-withdrawn-targets`) date from the first lists of the dataset, which were lists of drug targets. They apply to the genes of every list, and only the oncology lists have a registry date.

The registry date shows when a record became public, and not when the record first named the target or met the rules of the modality. The date from which a target counts is therefore either established by an archived registry copy or inferred (`known_from_basis, analysis/known_from_basis.tsv`). It is inferred when it falls on or before 2017-03-09, because the earliest archived copy cannot show that the record then qualified. This applies to 39 of the 65 ADC targets, 13 of 31 TCE targets and 59 of 76 IO targets. The other targets (26, 18 and 17) are established by an archived record or by a later trial start. CD5 has no registered trial and so no registry date, and it counts from the date of its 1991 publication. Dating the inferred targets no earlier than 2017-03-09 changes no label at any cutoff from 2017-03-31 on. At the three cutoffs of 2016, no oncology target can be verified, so those cutoffs rest on inferred dates (`analysis/known_from_basis_sensitivity.tsv`). The inflammatory bowel disease lists have no registry date. They are dated by `first_trial` under both settings (`trial_start_only`), so a replay with them establishes when a trial started and not when it became public.

#### Inflammatory bowel disease lists

The lists are chosen by indication. A gene is in a list when a drug that Open Targets lists as acting on it is listed for Crohn’s disease, ulcerative colitis or inflammatory bowel disease. The drug must also have an interventional study in that indication on ClinicalTrials.gov. A drug acts on a gene through its mechanism of action in Open Targets, and protein families of at most four members and protein complexes are expanded to their genes. The studies are found among the trials that the Open Targets clinical reports of the indication cite and by a search on the names of the drug. Each gene is dated by the first such study.

#### Lists of the claim association

These lists are not lists of drug targets. There are three kinds. (1) The GWAS lists hold the genes with a genome-wide significant variant in the GWAS Catalog [35] for type 2 diabetes, lipid traits, obesity, psoriasis and inflammatory bowel disease. A gene counts from the later of the publication and the Catalog entry of the first study that reports one (`.gwas`) or of the second, independent study (`.gwas-rep`, no look-ahead from a single study). (2) The cancer driver list holds the 633 driver genes of the IntOGen compendium [36]. A gene counts from the first of five releases (2014-12, 2019-11, 2020-02, 2023-05 and 2024-09) that lists it, and a gene that the latest release dropped is not in the list. (3) The ClinGen lists [37] hold the genes with a Definitive or Strong gene–disease validity for lipid traits (11 genes) and obesity (5 genes), dated by the classification date. That date is a curation milestone and not the date of discovery.

#### Dating and left-censoring

Every source starts at some date, and a gene that the first record of a source already shows has been known since then or earlier. This applies to 212 of the 633 cancer driver genes (33%), which are dated by the first IntOGen release (2014-12). It also applies to a drug tested before trials were registered (about 2000), which is dated by its first registered trial. The genes dated by the first record of IntOGen are known at every deposited cutoff, because the first cutoff is 2016-06-30. The censoring therefore creates no false later gene. It means that the age of these known genes is not known.

The field `date_basis` of a list names the basis of its dates (`trial_start, first_study, second_study, release or classification_date`). Only the oncology lists have a registry date. The features that are built from the evidence of a list itself are named for each kind in evidence kinds.json (`same_source_features`). They are the Open Targets genetic-association score for a GWAS list and the somatic-mutation score for the cancer drivers. They would give the answer away, and an evaluation of that list should leave them out.

Curated negatives and eligibility gates are defined per area (`areas/<area>/area.json`). The oncology negatives are genes that a rule marks as unlikely targets. There are three groups. Pan-essential genes are the common essentials of DepMap. Housekeeping genes are expressed at 10 nTPM or more in at least 90% of Human Protein Atlas tissues, with a median of at least 20 nTPM. Stromal or immune genes have a malignant specificity of at most *−*0.05 and a detection of at least 0.2 in non-malignant cells, in at least half of the cancers with single-cell data. A known gene is never a negative. The rules read the releases in force at the cutoff, so the negatives of a cutoff differ from today’s. The stromal or immune rule is skipped at cutoffs where its single-cell tables, which have no dated release, are not requested. The other three areas define no negatives. The eligibility gates are tables that flag genes expressed in the tissues or cells of the disease. In oncology they use tumour cohorts, DepMap lineage expression and single-cell malignant specificity. In inflammatory bowel disease they use gut and immune-cell expression, in psoriasis skin and immune-cell expression, and in metabolism the expression in the tissues of metabolic disease. The gates are annotations that a user may apply as a filter, and the label tables are not filtered by them.

### 2.10 Citations of the known genes

A known gene can be traced to the documents that made it known. The table `literature/gene_citations.tsv` has one row per gene, list and citation (a PubMed identifier or a ClinicalTrials.gov number) with the date from which the document was public (`herald-data citations`; 61,815 rows, 725 papers, 1,135 trials, 6,503 genes). For the GWAS lists the citations are the studies of the GWAS Catalog that report a genome-wide significant variant of the gene for the disease (560 studies), dated like the list by the later of the publication and the Catalog entry, so that the first citation of a .gwas list carries the date of the list. For the inflammatory bowel disease, ADC and TCE lists they are the interventional studies of the drugs that put the gene on the list, dated by the later of the start and the first submission to the registry (AACT copy of 2026-09-01); the IO list keeps only the study that dated the gene. The 165 papers of the oncology curation, with their claim types, are included at the resolution of the year. The lists of ClinGen and IntOGen have no rows, because their sources give labels, curation records and cohorts instead of papers. Rows dated after the newest release of the AACT copy and the Catalog are left out (planned start dates, papers dated ahead of print). Only identifiers and dates are released, and the sentences of the papers are not. The table is meant for the evaluation of methods that read documents. It is not a feature, and no ranking of the dataset reads it.

### 2.11 Audit of the automatic labels

The audit covers the lists dated by clinical trials (the oncology and inflammatory bowel disease areas), whose members and dates come from matching text records. The other lists take their members and dates from the records of their source. The dates of the GWAS lists are compared with PubMed (Technical Validation, Dates of the GWAS lists). The precision of the automatic matching was estimated on a random sample (`herald-data labels sample`, 50 targets per area, seed 1) of the targets that no manual audit decided. Each gene–drug–trial record of the sample was assessed in two steps, first by an automated review and then by a human reviewer.

The automated review is an AI-generated workflow with programmed logic. For each record it assessed the proposed association from the trial-registry record and from external biomedical databases (ChEMBL, DrugBank through UniProt, and PubMed), and it returned a categorical judgement, a rationale, a verbatim quotation of the evidence and its source. The databases serve the audit only. They are not inputs of the deposit. The first step assessed the indication in the inflammatory bowel disease area and the modality in the oncology area. The indication assessment considered the disease context, the intervention and the outcome measures. A disease-relevant primary outcome supported a positive judgement at high confidence. When no primary outcome matched, a relevant secondary outcome supported it at medium confidence, following the reporting roles of outcomes [38]. A registered outcome shows that the indication was investigated, not that the intervention was beneficial. The modality (ADC or T-cell engager) was identified in the trial record from nomenclature patterns and selected molecule names. The second step assessed the relationship between the gene and the drug, in the trial record or in the external databases, by retrieving terms of the mechanism of the drug and the gene. A separate evidence check retrieved each cited source and compared it with the quotation by three-token n-gram overlap [39]. The outputs of every record are saved as JSON.

A human reviewer (a co-author, Y.W.) judged every record of the sample (100 of 100) and entered a verdict (correct, incorrect or unclear) and a note. For each record the reviewer assessed the indication and the target of the drug in the inflammatory bowel disease area, and the indication, the target and the drug modality in the oncology area. A positive indication match meant that the drug was an intervention associated with a change in a primary or secondary outcome for the disease of interest; conditioning medications were excluded. A positive target match meant that the gene or its protein was a direct target of the drug, and a positive modality match that the drug was an ADC or a T-cell engager. The reviewer used the trial record, ChEMBL, DrugBank and PubMed. A record was correct when every criterion was positive, and incorrect when any was negative. A record with a partial match and an undetermined remainder was unclear. The reviewer did not see the verdict of the automated review, so the judgement was blinded to it. Verdicts of the automated review and of the reviewer were compared on the 98 records that both judged correct or incorrect. A record for which either verdict was unclear was reported separately and left out of the confusion matrix. We report agreement, Cohen’s kappa, accuracy, precision, recall, specificity, the F1 score and the negative predictive value, with the confidence of the automated review, the step that failed in each disagreement, and the notes of the reviewer, which we grouped by theme. The precision of the labels and its Wilson interval were computed from the verdicts of the reviewer (`herald-data labels score`).

### 2.12 Identifier harmonization

Gene symbols are harmonized to the gene universe of each disease area at build time. A per-area crosswalk table in the deposit maps symbols to UniProt, Ensembl and Entrez accessions. The crosswalk is keyed on gene symbol. It is assembled from sources that carry several identifiers in one row. These are the Human Protein Atlas gene table (symbol, Ensembl and UniProt, with the first accession when a row lists several), the DepMap column headers (symbol and Entrez), the UniProt fallback of the area and the Entrez identifiers of the curated literature table. It is built once from the working copy of the release archive and is not rebuilt per cutoff. No separate table of retired or merged identifiers is used, and a symbol without an accession in any of these sources is left empty. Such genes stay in every table with empty accession fields and are not excluded from any analysis or label. The gene universe (the protein-coding genes of the current Human Protein Atlas) and the crosswalk are not rebuilt per cutoff. Some hindsight therefore remains. A gene that was named or annotated after a cutoff is present at it, with the features that its releases supply (Table 7 shows how many genes have data at each cutoff).

**Table 7:** Coverage of the features at three year-end cutoffs. Features: full set (biology-only set in parentheses); genes: genes with at least one feature, of the 20,151 genes of the universe.

| Area | Features 2016 | Features 2020 | Features 2024 | Genes 2016 | Genes 2020 | Genes 2024 |
|---|---|---|---|---|---|---|
| Oncology | 30 (20) | 57 (36) | 68 (44) | 18,750 | 19,600 | 20,151 |
| IBD | 30 (20) | 58 (36) | 66 (41) | 18,750 | 19,600 | 20,151 |
| Metabolism | 30 (20) | 58 (36) | 66 (41) | 18,750 | 19,600 | 20,151 |
| Psoriasis | 30 (20) | 58 (36) | 66 (41) | 18,750 | 19,600 | 20,151 |

### 2.13 Ethics statement

All inputs are public, aggregate-level resources: no human participants were recruited, no individual-level data are included, and no access-controlled source is used.

### 2.14 Use of AI tools

Claude Haiku/Sonnet/Opus 5.5 were used to assist with software development, testing, and manuscript preparation. All AI-assisted code and manuscript content were reviewed and validated by the author, who takes full responsibility for the work.

## 3 Data Records

The dataset is deposited in Zenodo [40]. It is distributed as the public tree of the versioned herald-data package, a self-contained directory that is copyable as a whole and that carries its own provenance manifest. Its layout is:

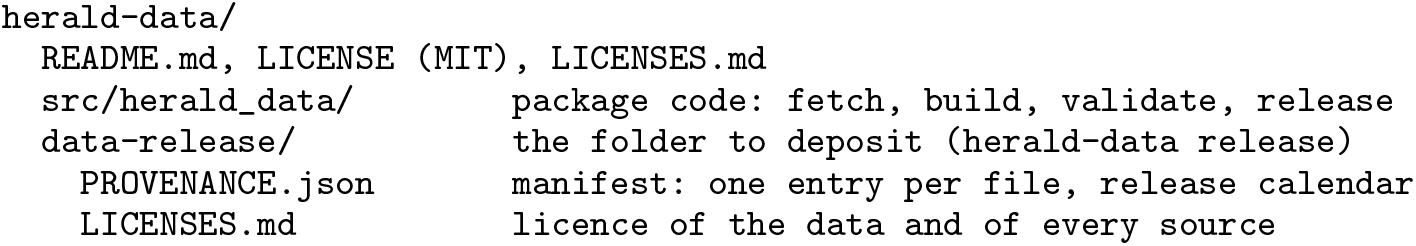
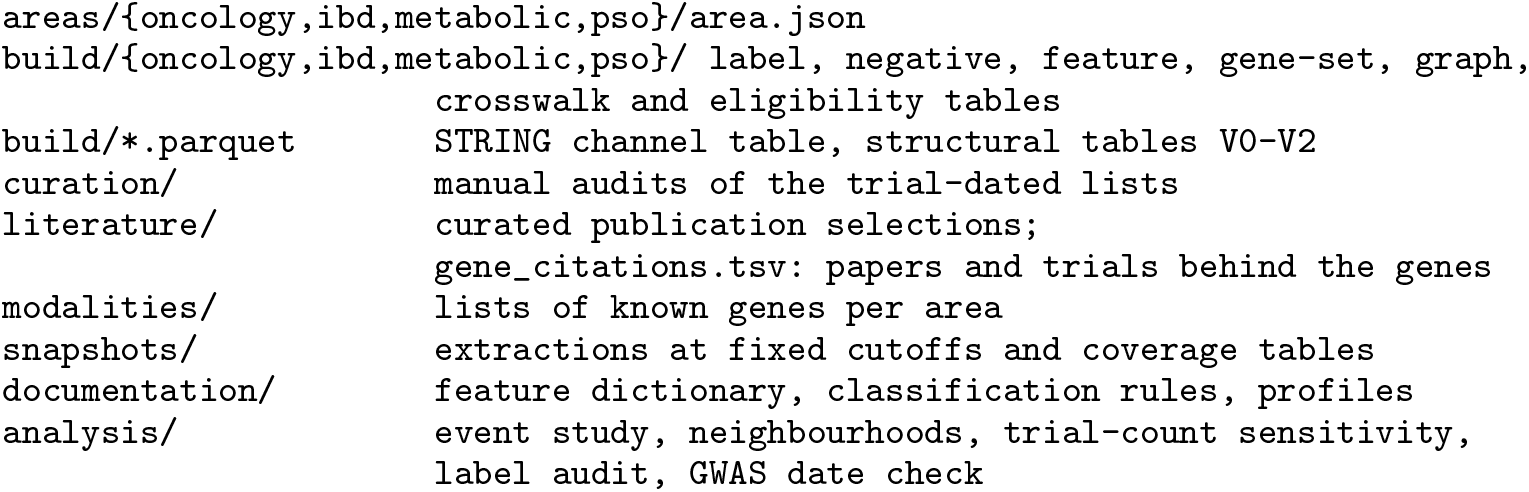

Table 9 describes each file type and Table 10 defines the fields of the released tables. The manifest (`PROVENANCE.json`) is generated from the deposit. For every file it records the size, a checksum and the licence, together with the notices that a source asks for (for example that of the GTEx Portal). The manifest also contains the release calendar of the source archive. For every archived release of every source, the calendar gives the release date, the evidence for that date, the files and bytes of the release, and the access date. A second calendar covers the supplementary sources that are archived for reuse. Both calendars are JSON files that the accompanying code generates (Usage Notes). The upstream release archives `(raw/<sor>/<release>/`, gigabytes of downloads) are not part of the deposit. The same holds for the working copies that the build derives from them: the converted ClinicalTrials.gov registry copies, the converted Open Targets releases and the structural tables of each AlphaFold DB release. The release calendar records the archive, and `herald-data fetch` followed by `herald-data build` regenerates both. The per-area feature registry gives the provenance class, risk grades and availability date of every feature (`herald-data registry`). The deposit also holds the feature dictionary, the classification rule table (`documentation/`) and the extractions at fixed cutoffs with their coverage tables (`snapshots/`).

**Table 8:** The 19 lists of known genes. The claim is *target* when a drug acting on the gene was tested and *association* when the gene is linked to a disease or trait. The scope is the kind of drug (modality) or the disease. Genes: size of the list. P/F: genes known (P) and still to become known (F) at the end of the year, under the default dating (the later of the first trial and the registry date for the oncology lists; the date of the list for the others).

| Area | List | Claim | Scope | Dated by | Genes | P/F 2016 | P/F 2020 | P/F 2024 |
|---|---|---|---|---|---|---|---|---|
| Oncology | ADC.trial | target | modality | first registered trial | 65 | 39/26 | 52/13 | 61/4 |
| Oncology | CANCER.driver | association | disease | first IntOGen release | 633 | 212/421 | 502/131 | 633/0 |
| Oncology | IO.trial | target | modality | first registered trial | 76 | 57/19 | 74/2 | 76/0 |
| Oncology | TCE.trial | target | modality | first registered trial | 31 | 13/18 | 21/10 | 28/3 |
| IBD | CD.trial | target | disease | first registered trial | 93 | 73/20 | 83/10 | 89/4 |
| IBD | IBD.gwas-rep | association | disease | second GWAS study | 233 | 101/132 | 127/106 | 213/20 |
| IBD | IBD.gwas | association | disease | first GWAS study | 483 | 180/303 | 250/233 | 418/65 |
| IBD | IBD.trial | target | disease | first registered trial | 136 | 93/43 | 117/19 | 130/6 |
| IBD | UC.trial | target | disease | first registered trial | 99 | 64/35 | 81/18 | 94/5 |
| Metabolism | LIPIDS.clingen | association | disease | classification date | 11 | 0/11 | 3/8 | 11/0 |
| Metabolism | LIPIDS.gwas-rep | association | disease | second GWAS study | 2,091 | 136/1,955 | 606/1,485 | 1,580/511 |
| Metabolism | LIPIDS.gwas | association | disease | first GWAS study | 3,145 | 244/2,901 | 1,196/1,949 | 2,459/686 |
| Metabolism | OBESITY.clingen | association | disease | classification date | 5 | 0/5 | 2/3 | 4/1 |
| Metabolism | OBESITY.gwas-rep | association | disease | second GWAS study | 2,482 | 37/2,445 | 1,288/1,194 | 2,332/150 |
| Metabolism | OBESITY.gwas | association | disease | first GWAS study | 3,861 | 166/3,695 | 2,220/1,641 | 3,590/271 |
| Metabolism | T2D.gwas-rep | association | disease | second GWAS study | 608 | 17/591 | 297/311 | 438/170 |
| Metabolism | T2D.gwas | association | disease | first GWAS study | 1,416 | 64/1,352 | 619/797 | 1,063/353 |
| Psoriasis | PSO.gwas-rep | association | disease | second GWAS study | 102 | 23/79 | 43/59 | 53/49 |
| Psoriasis | PSO.gwas | association | disease | first GWAS study | 357 | 61/296 | 198/159 | 246/111 |

**Table 9:** Files in the deposited dataset (paths relative to data-release/; <area> is oncology, ibd, metabolic or pso).

| Path | Format | Content |
|---|---|---|
| PROVENANCE.json | JSON | generated manifest: one entry per deposited file, and the release calendar of the source archive (Table 10) |
| areas/<area>/area.json | JSON | declarative definition of the disease area: diseases, gene universe, labels, eligibility gate |
| build/<area>/labels_pnu_<area>.parquet | Parquet | per-gene positive-unlabeled labels (Table 10) |
| build/<area>/negatives_<area>.parquet | Parquet | curated negative pool with selection reasons |
| build/<area>/crosswalk_<area>.parquet | Parquet | gene symbol to UniProt, Ensembl and Entrez accessions |
| build/<area>/gene_leaves_opentargets_<area>.parquet | Parquet | per-gene feature table (one column per feature) |
| build/<area>/leaf_meta_*.parquet | Parquet | one row per per-gene feature leaf, with source and role |
| build/<area>/gene_sets_*.json | JSON | frozen gene sets for the over-representation view (KG entities + pathways) |
| build/<area>/graph_components_*.parquet | Parquet | graph context components of the knowledge-graph view |
| build/<area>/elig/<disease>.parquet | Parquet | eligibility gate per disease |
| build/<area>/gwas_annotation_<area>.parquet | Parquet | GWAS Catalog association table (annotation only, not a feature) |
| build/oncology/registry_dates_oncology.tsv | TSV | registry date, with its delay in days from the first trial, of each oncology gene (Table 10) |
| build/<area>/string_ppi_edges.parquet | Parquet | high-confidence protein–protein-interaction edges (gene, neighbour) |
| build/string_v12_detailed.parquet | Parquet | STRING v12 per-channel scores |
| build/structural_v*.parquet | Parquet | per-gene structure-derived features |
| curation/*.tsv | TSV | manual audits of the trial-dated lists (97 records, two tables), the public drug records behind 19 ADC targets (19 rows) and the hand-reviewed immuno-oncology drug table (76 genes) |
| literature/ | TSV, Parquet, JSON | curated publication selections and their fetch manifest; the dated citations (PMID, NCT) behind the known genes of each list |
| modalities/<area>/<list>.tsv | TSV | lists of known genes (Table 10) |
| snapshots/<area>/<cutoff>/ | Parquet, TSV, JSON | features of the full set, their registry, the P/F/D/N/U label of every list and the releases used, at 36 cutoffs (quarter ends 2016-06-30 to 2024-12-31 and 2020-02-03) |
| snapshots/coverage_*.tsv | TSV | features and genes with data per cutoff and feature set; P/F/D/N/U per cutoff and list under both dating rules |
| documentation/*.tsv | TSV | feature dictionary, classification rule table and risk ceilings of the feature sets |
| analysis/*.tsv, analysis/event_study.pdf, analysis/event_study_curves.pdf, analysis/neighbours.pdf | TSV, PDF | event study and neighbourhood analyses of Technical Validation, with the trial-count sensitivity of the lists, the label audit (label_audit_*.tsv, label_precision.tsv) and the release-date check (release_date_check.tsv, release_date_audit_sample.tsv) |

**Table 10:**
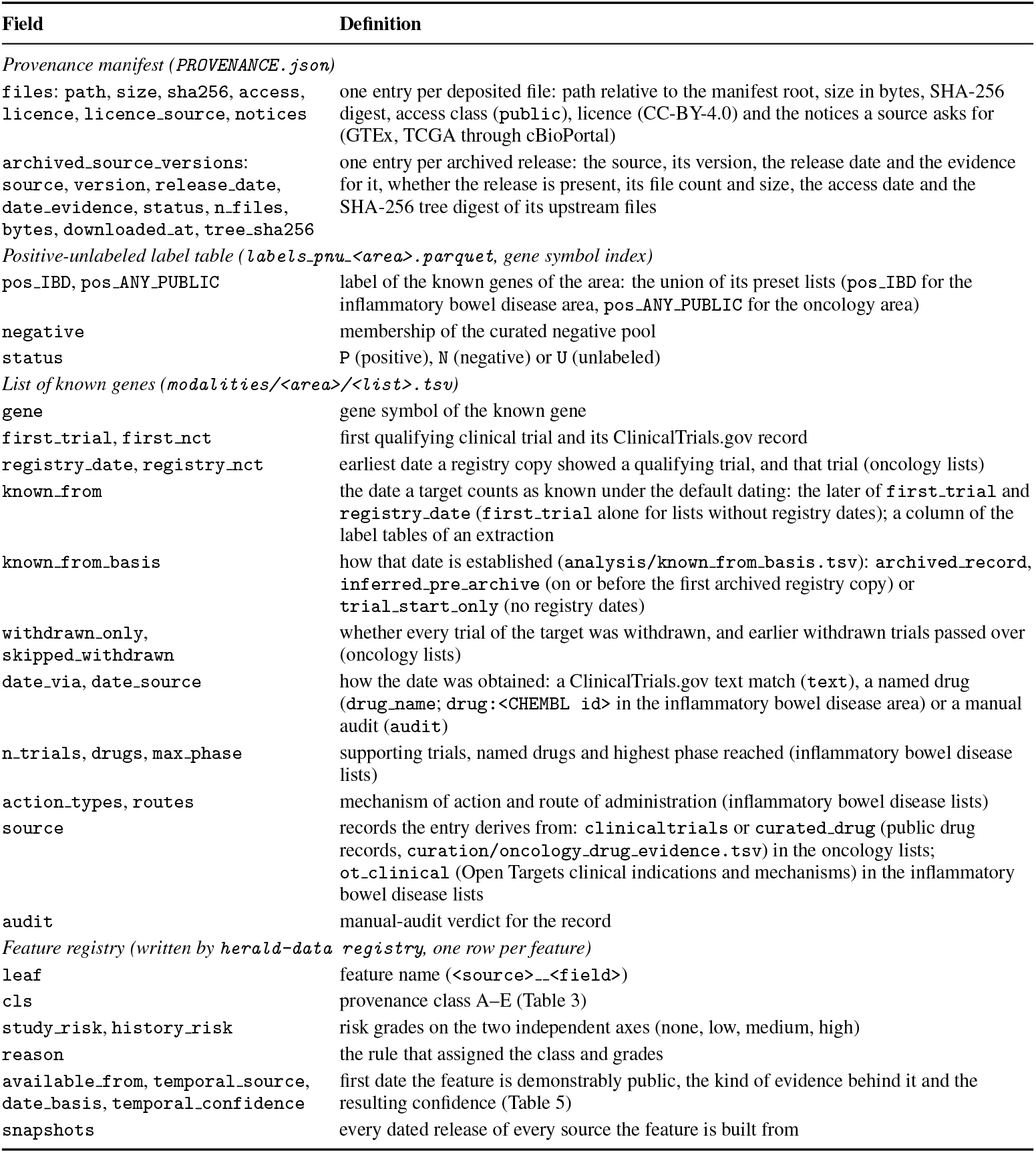
Fields of the released tabular records.

| Field | Definition |
|---|---|
| *Provenance manifest (PROVENANCE.json)* |  |
| files: path, size, sha256, access, licence, licence_source, notices | one entry per deposited file: path relative to the manifest root, size in bytes, SHA-256 digest, access class (public), licence (CC-BY-4.0) and the notices a source asks for (GTEx, TCGA through cBioPortal) |
| archived_source_versions: source, version, release_date, date_evidence, status, n_files, bytes, downloaded_at, tree_sha256 | one entry per archived release: the source, its version, the release date and the evidence for it, whether the release is present, its file count and size, the access date and the SHA-256 tree digest of its upstream files |
| *Positive-unlabeled label table (labels_pnu_<area>.parquet, gene symbol index)* |  |
| pos_IBD, pos_ANY_PUBLIC | label of the known genes of the area: the union of its preset lists (pos_IBD for the inflammatory bowel disease area, pos_ANY_PUBLIC for the oncology area) |
| negative | membership of the curated negative pool |
| status | P (positive), N (negative) or U (unlabeled) |
| *List of known genes (modalities/<area>/<list>.tsv)* |  |
| gene | gene symbol of the known gene |
| first_trial, first_nct | first qualifying clinical trial and its ClinicalTrials.gov record |
| registry_date, registry_nct | earliest date a registry copy showed a qualifying trial, and that trial (oncology lists) |
| known_from | the date a target counts as known under the default dating: the later of first_trial and registry_date (first_trial alone for lists without registry dates); a column of the label tables of an extraction |
| known_from_basis | how that date is established (analysis/known_from_basis.tsv): archived_record, inferred_pre_archive (on or before the first archived registry copy) or trial_start_only (no registry dates) |
| withdrawn_only, skipped_withdrawn | whether every trial of the target was withdrawn, and earlier withdrawn trials passed over (oncology lists) |
| date_via, date_source | how the date was obtained: a ClinicalTrials.gov text match (text), a named drug (drug_name; drug:<CHEMBL id> in the inflammatory bowel disease area) or a manual audit (audit) |
| n_trials, drugs, max_phase | supporting trials, named drugs and highest phase reached (inflammatory bowel disease lists) |
| action_types, routes | mechanism of action and route of administration (inflammatory bowel disease lists) |
| source | records the entry derives from: clinicaltrials or curated_drug (public drug records, curation/oncology_drug_evidence.tsv) in the oncology lists; ot_clinical (Open Targets clinical indications and mechanisms) in the inflammatory bowel disease lists |
| audit | manual-audit verdict for the record |
| *Feature registry (written by* herald-data *registry, one row per feature)* |  |
| leaf | feature name (<source>__<field>) |
| cls | provenance class A–E (Table 3) |
| study_risk, history_risk | risk grades on the two independent axes (none, low, medium, high) |
| reason | the rule that assigned the class and grades |
| available_from, temporal_source, date_basis, temporal_confidence | first date the feature is demonstrably public, the kind of evidence behind it and the resulting confidence (Table 5) |
| snapshots | every dated release of every source the feature is built from |

The tables are released under CC BY 4.0 and the code under the MIT licence. The `herald-data release command` removes what the dataset licence cannot carry. These are the sentences quoted from the papers in the literature table, every KEGG gene set (restricted by MSigDB), and the short-lived-protein features of Li et al. [41] (an Elsevier user licence, not a Creative Commons licence). Two features count, for each gene, the KEGG pathway sets that contain it; the sets themselves are not deposited. The inflammatory bowel disease trial lists are built from the Open Targets Platform alone, the oncology lists rest on ClinicalTrials.gov records and public drug records, and the association lists on the records of their sources (GWAS Catalog, ClinGen, IntOGen) with the dates of those records. The manifest gives the licence of each file and the notices that a source asks for, and `LICENSES.md` lists the licence of every source (Table 2). No ChEMBL content is deposited: ChEMBL is licensed CC BY-SA 3.0, which the dataset licence cannot carry, so the deposited oncology lists have no molecule columns and the drug tables built from ChEMBL are not shipped. The release step checks its own result against the licences (the identifiers and columns of ChEMBL, the names of restricted and proprietary sources, restricted features and gene sets, quoted sentences) and stops when it finds anything. The folder passes `herald-data validate`.

## 4 Data Overview

Figure 2 and Table 1 show what an extraction provides on 2020-02-03, the release date of DepMap 20Q1, for one list of each area. The extractions hold 47 to 53 features of the full set for 20,151 genes. Each gene is labelled for a list as known on or before the cutoff (P) or known after it (F), and the genes are placed by a UMAP of the features of the area. The labels play no part in the placement. In the ADC list the 49 genes labelled P lie in a narrow part of the feature space, and most of the 16 genes labelled F lie in or next to it, with a few apart. The lists differ in size, from 16 genes labelled F in the ADC list to 498 in the second-study list of type 2 diabetes. A method that learns from the genes labelled P can be tested on how highly it ranks the genes labelled F. The ADC list is compared with surface features in Technical Validation (Fig. 6).

**Figure 2:**
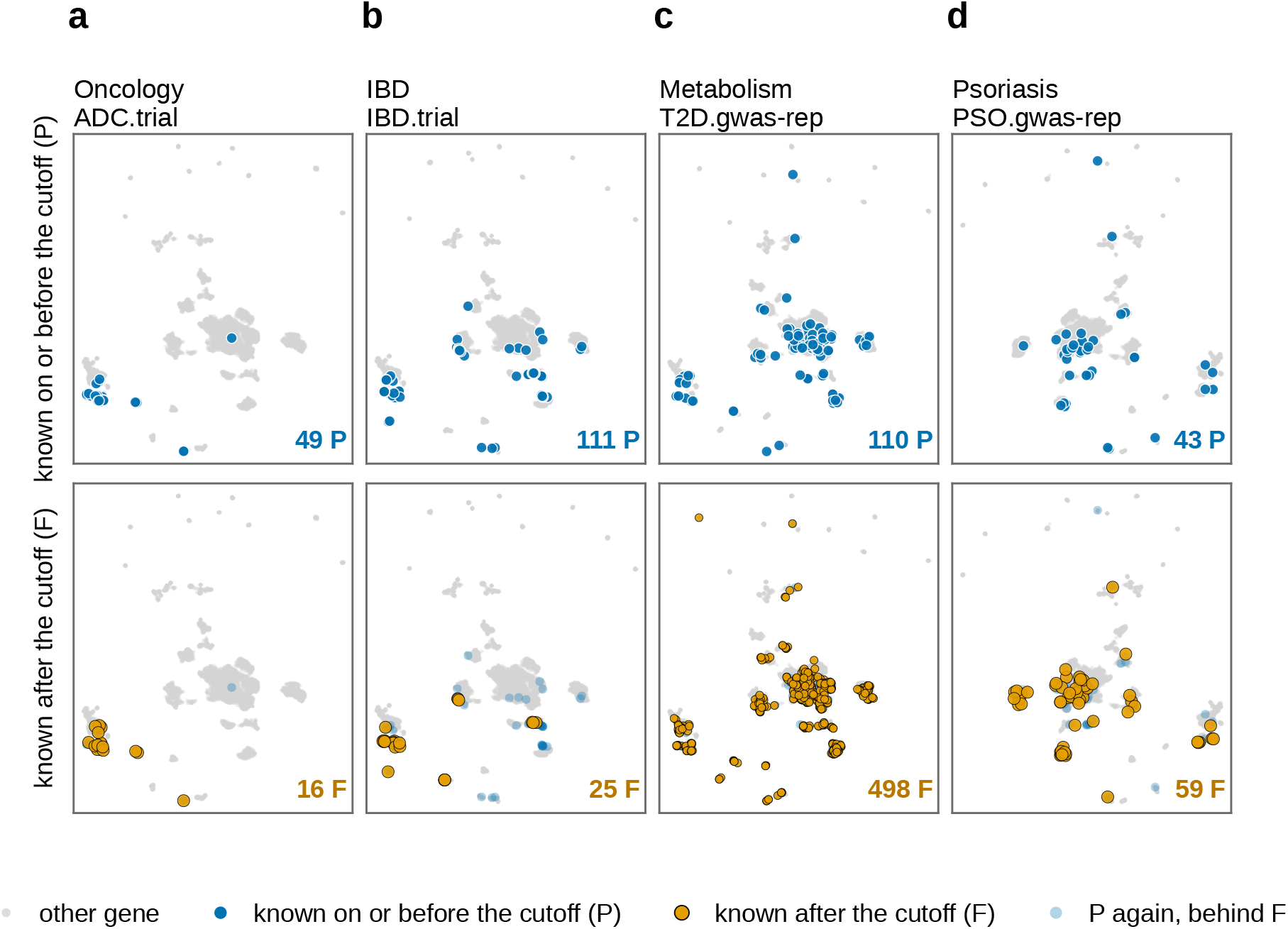
Labels of one list of each area at the cutoff 2020-02-03 in the feature space of that date. UMAP of 20,151 genes by the features of the full set of the area (Table 1); the labels were not used to place the genes, and the gene positions are the same in the upper and the lower row of a column. Columns: the ADC list of oncology (**a**), the list of the targets of drugs tested in inflammatory bowel disease (**b**), the second-study GWAS list of type 2 diabetes in metabolism (**c**) and the second-study GWAS list of psoriasis (**d**). Upper row: genes known on or before the cutoff (P, blue). Lower row: genes known after it (F, orange), with the P genes faint behind them. The numbers of P and F genes are given in the panels. Genes of both kinds often lie within the same groups.

## 5 Technical Validation

### 5.1 Temporal admissibility

The release calendar gives the release date of every archived release together with its evidence, and the resolver selects only releases dated on or before the cutoff. A record that is later than the cutoff therefore cannot enter a snapshot. Each extraction records the release resolved for every source (`releases in extract.json`), and source releases that postdate the cutoff are excluded from that snapshot. Package tests check that a feature is excluded at a cutoff before the first release of its source and admitted after it. For example, no DepMap feature is admitted on the day before DepMap 20Q1, and no AlphaFold feature at 2020-02-03.

### 5.2 Leakage segregation

The provenance class and the two risk grades of every feature are written by `herald-data registry` (`feature_registry.tsv`). They are assigned from the source and the feature name by a rule table that is released with the code. The feature sets of Table 4 follow from the grades by thresholding alone, so the components that a feature set admits can be verified without running any model. The labels of every area are built only from the released lists of known genes, so no score or list that is not public enters the labels or the features. `herald-data` validate fails when such a file is found in a public folder. Package tests check three properties. The class of each feature matches the rule table. The feature set is applied before the class filter, so a class-E feature stays out of the biology-only set even when class E is requested. The class filter keeps only the requested classes.

### 5.3 Example of a resolved cutoff

The resolver can be checked on a single cutoff. For the inflammatory bowel disease area, with Open Targets as the knowledge graph and a cutoff of 2022-02-08, it selected Open Targets 21.11, UniProt 2021 04, Human Protein Atlas v20.1 (blood proteome v20), GTEx V8, DepMap 22Q1, MSigDB 7.5.1, STRING v11.5 and gnomAD v2.1.1. The full feature set then held 52 features (31 of class A, 1 of B, 15 of C, 4 of D and 1 of E) for 20,151 genes, and the biology-only set the 31 class-A features. At the end of 2024 the full set holds 66 features. The extraction was run on the build of the deposit, which leaves out the short-lived-protein feature of Li et al.

### 5.4 Release calendar

The release calendar of the source archive lists 183 archived releases of 20 sources (Fig. 3). Every release carries a release date and the evidence for it, as a text field of the manifest (`date_evidence`). The field holds the changelog line, the release-table row, the file or folder timestamp, the publication or the version name that was used. The evidence is a release record for 82 releases, a file or folder timestamp for 70, a version name for 25 and a publication date for 6 (Table 5). Three further entries have the status `current-moved` and are not archived releases. A second calendar lists the 54 supplementary releases of six further sources, including the 39 AACT copies, with their dates, date evidence and licences. `herald-data validate` fails when a supplementary release is recorded as present without a date, without date evidence or without its files on disk.

**Figure 3:**
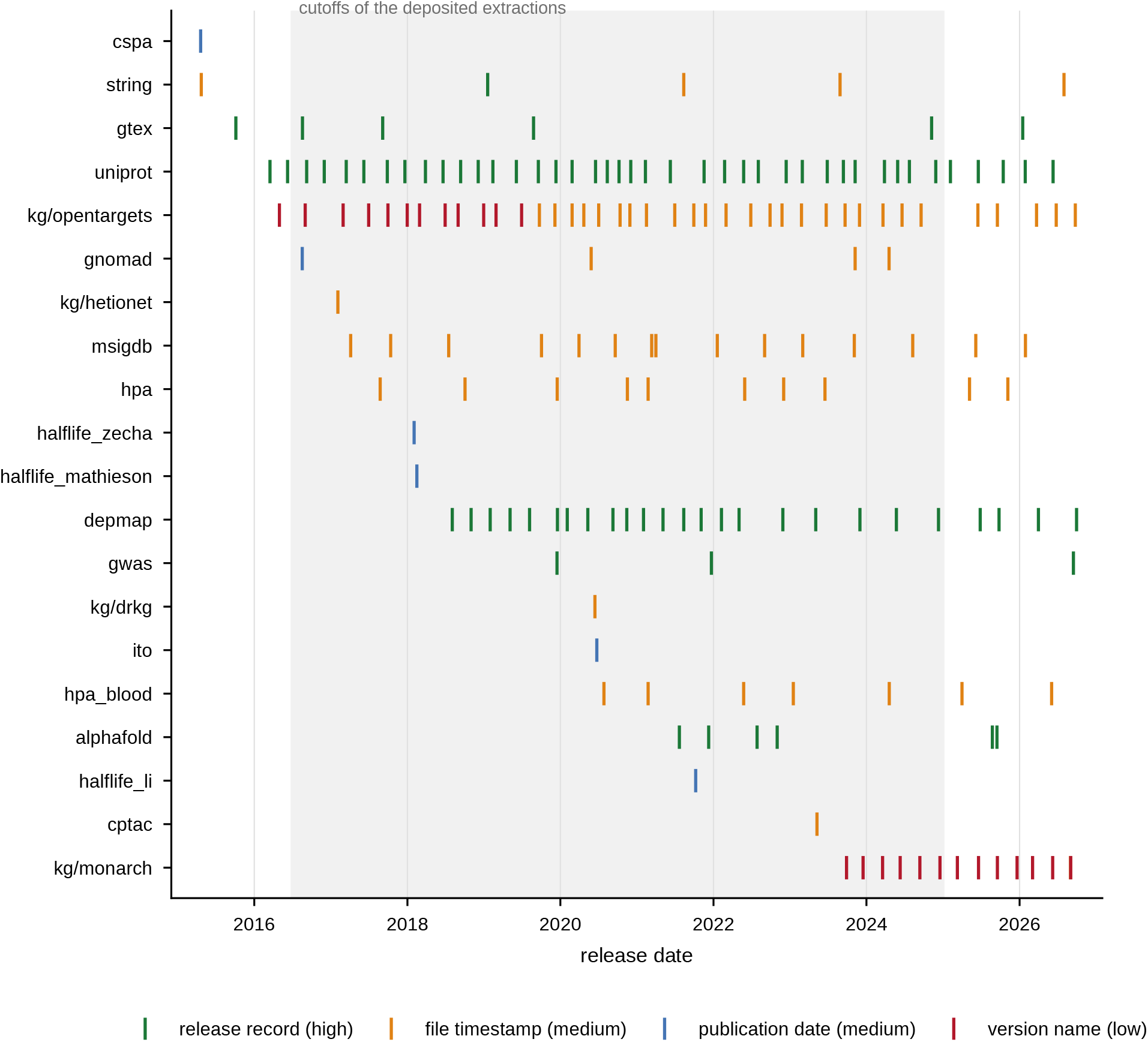
Release timeline of the sources. Each tick is an archived release, coloured by the kind of evidence that dates it. The shading covers the quarter-end cutoffs of the deposited extractions (2016-06-30 to 2024-12-31). A cutoff can use a source only from its first release onward, and the number of features and genes with data at each cutoff follows from it (Table 7).

The dates are release-level: the dataset carries no per-record dates for the evidence inside a release. A date that rests on a file timestamp or a version name is a proxy for the day the release became public. The dates have been compared with independent records for a sample only (`analysis/release_date check.tsv`). The sample was drawn at random within each kind of date evidence (36 releases, `analysis/release_date_audit_sample.tsv`), and 27 of the 36 were compared, 26 of them to the day. The calendar date agreed with the independent date for 9 releases, was earlier for 9 and was later for 8. Six of the 27 are dated by the publication of the paper that released the data, and in all six the date is the electronic publication date that PubMed (or Crossref) gives. Nine are dated by a file timestamp. For six of them the calendar date is earlier than the date of the provider’s release page, announcement or release notes (Human Protein Atlas versions 19 and 20, gnomAD v4.1, Open Targets 19.09 and a Monarch build, each by 1 or 2 days, and STRING v12.5 by 59 days, because its links file was modified on 1 August 2026 and the provider dates the version from 29 September 2026). The DRKG archive is dated 52 days later than the first commit of its repository, Hetionet agrees, and for MSigDB the release notes give the month only. Nine are releases dated by the name of a folder or a version. Of the four Monarch releases, two agree with the date of the GitHub release of the software and the folder date is 11 and 29 days earlier for the other two; the five Open Targets releases are dated by the end of the month of their version name, 1 to 14 days later than the post of the release in the project blog. Of ten releases dated by a release record of the provider, three have a second record: the calendar date is 23 days earlier than the announcement of AlphaFold DB version 6 (the changelog dates the files), and 10 and 1 days later than the DepMap 23Q4 announcement and the creation of the GTEx v11 files. Where the calendar date is earlier, a cutoff that falls in the interval would admit a release that was not yet public, if the files were not public from the earlier date. Of the 9 earlier dates, 5 are within 2 days, and the others are 11, 23, 29 and 59 days (STRING v12.5 matters only for cutoffs after August 2026). The other 9 releases of the sample were not compared, because no independent dated record was found or the only second record is the provider’s own (the searches are in the sheet). The availability of a feature is the first dated release of each source it is built from. It is not checked field by field, with one exception: gnomAD’s first release, ExAC, has no loss-of-function observed/expected ratio, so that feature is dated from v2.1.1. A feature that an early release supports only in part has missing values in that snapshot, which the counts of genes with data show (Table 7).

### 5.5 Behaviour of the features under the two axes

Two descriptive analyses of the deposited extractions show how the features behave under the time and leakage axes (Fig. 4 and Fig. 5). Neither fits a model. Both are computed from the `snapshots/` tables alone (`herald-data leakage` and `herald-data` neighbours; results in `analysis/`).

**Figure 4:**
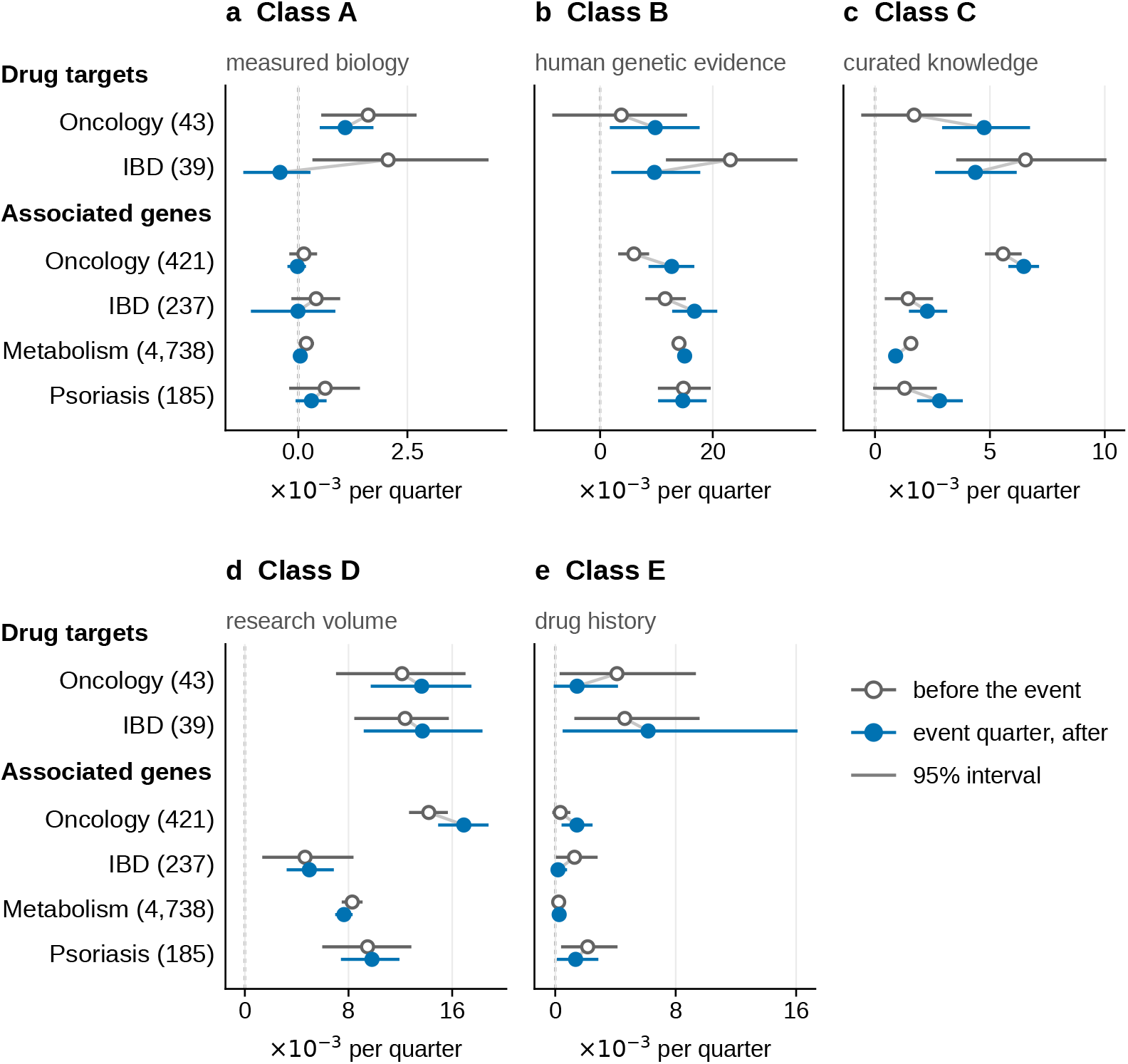
Event study of the features under the time and leakage axes, computed from the deposited extractions. Each panel is one provenance class (**a**, class A, measured biology; **b**, class B, human genetic evidence, the single feature Open Targets genetic association; **c**, class C, curated knowledge; **d**, class D, research volume; **e**, class E, drug history, the single feature Open Targets known drug). Each row is a group of genes that became known in a list of an area, either as drug targets (the trial lists of oncology and inflammatory bowel disease) or as associated with a disease (the oncology drivers and the GWAS lists of inflammatory bowel disease, metabolism and psoriasis). The number of genes with quarters on both sides of the event is in brackets. Circles show the mean excess quarterly change of the rank percentile, in units of 10*^−^*^3^ per quarter, over the eight quarters before the event quarter (open) and from the event quarter on (filled). The excess is relative to control genes that started the quarter in the same decile. Horizontal lines are 95% bootstrap intervals over genes. The curves of every quarter are in the deposited analysis folder.

**Figure 5:**
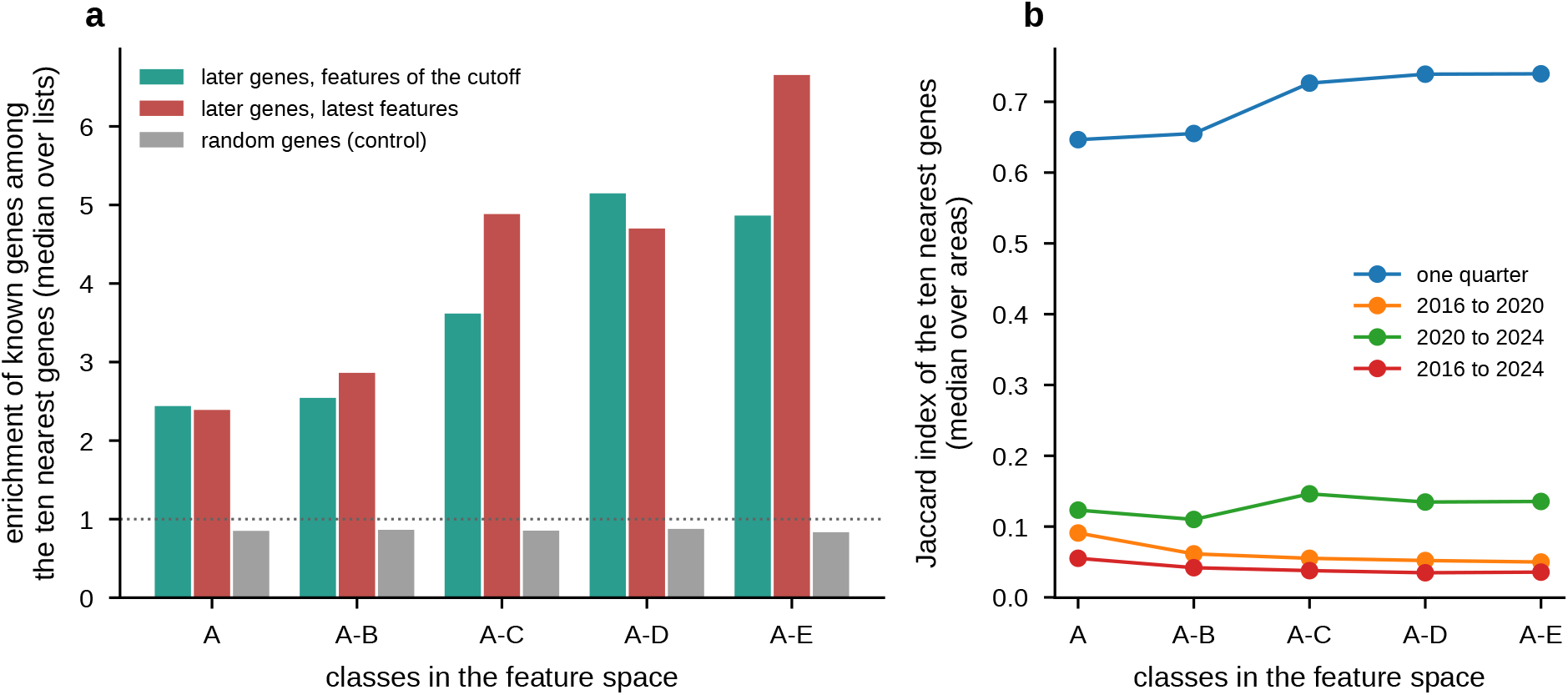
Neighbourhoods of the genes in feature space. **a**, Enrichment of known genes among the ten nearest genes of the genes that were still to become known (later genes), with the features of the cutoff and with the latest features, and of random genes in no list (median over 90 list–cutoff pairs), for five feature sets (classes A, A to B, A to C, A to D and A to E). A value of 1 is the share of known genes among all genes. **b**, Jaccard index of the ten nearest genes of a gene at two snapshots, one quarter apart and four or eight years apart (median over the four areas).

#### Event study: design

The event of a gene is the quarter in which it became known in a list of its area, dated as the list dates it. The lists of each area were grouped by their claim, so that genes that became known as drug targets (the trial lists) and genes that became known as associated with a disease (GWAS, ClinGen and IntOGen) are followed separately. This gives six groups of an area and a claim (Fig. 4). For a GWAS disease the event is the first study. Genes that became known between 2016-07-01 and 2024-12-31 were used. The target groups hold 45 genes in oncology and 39 in inflammatory bowel disease, and the association groups hold 421 genes in oncology, 286 in inflammatory bowel disease, 4,805 in metabolism and 200 in psoriasis. For every feature, the genes were ranked at each quarter-end snapshot, and the rank percentile (0 to 1) was followed for eight quarters before and eight quarters after the event quarter. The change of the percentile over one quarter was compared with that of control genes. Control genes are in no list in any quarter and started the quarter in the same decile of that feature. The mean change of the controls was subtracted, so that regression to the mean and releases that move every gene cancel out. A gene whose window is cut by the first or the last snapshot contributes the quarters it has, and no quarter is imputed. Only features that exist and vary in every snapshot were used, so that a feature that appears in a later release is not read as a shift. There are 25 such features in each area (15 of class A, 1 of B, 6 of C, 2 of D and 1 of E; `analysis/event_study_features.tsv`). The results concern these features and not those introduced in later releases. Fig. 4 summarises each class and group by two numbers, the mean excess change over the quarters before the event quarter and the mean over the quarters from it on, each taken per gene first and then over the genes that have quarters on both sides of the event (95% bootstrap intervals over genes). The curves of every quarter are in `analysis/event_study curves.pdf`.

#### Event study: observations

For class A features, the mean excess change differs by at most 0.0025 percentile per quarter between the quarters before the event and those from the event quarter on, in all six groups. The largest difference (*−*0.0025) is a decrease, in the target genes of inflammatory bowel disease, and it is the only one whose interval excludes zero; every other difference is 0.0005 percentile or less in size. The two class D features record research volume. They change faster than the controls in every group, before the event as well as after it (0.005 to 0.017 percentile per quarter on average). The only group in which the interval of the difference excludes zero is the associated genes of oncology (0.014 before, 0.017 after). In the other five groups the difference is 0.002 or less in size, with intervals that include zero. The six class C features, curated knowledge, change by 0.001 to 0.007 per quarter. The interval of the difference excludes zero in three groups, with an increase in the target genes of oncology (0.002 before, 0.005 after) and the associated genes of oncology (0.006, 0.006), and a decrease in metabolism (0.002, 0.001). Class B is one feature, the Open Targets genetic-association score, which is built from the kind of evidence that dates the GWAS and ClinGen lists. In the associated genes of oncology its mean excess is higher after the event than before, with an interval that excludes zero (0.006 to 0.013). In inflammatory bowel disease the change is of a similar size (0.012 to 0.017) but its interval reaches *−*0.0002. In metabolism and psoriasis it is not seen (0.014 to 0.015, and 0.015 to 0.015), so the step is not seen for every list of its kind. In the target genes the intervals of the difference include zero and are wide (for example *−*0.008 to 0.021 in oncology). The one class E feature, Open Targets known drug, is near zero in the association groups (0.000 to 0.002; in oncology it rises from 0.0003 to 0.0014, an interval of 0.0001 to 0.0022 for the difference). In the target groups all intervals of the difference include zero, and the curves are flat except for isolated jumps of a few genes. The largest jump at the event quarter (inflammatory bowel disease, mean 0.025, 95% interval *−*0.0004 to 0.061) is matched by one of similar size in a quarter unrelated to the event (0.020 seven quarters before it; in oncology the largest, 0.030, falls six quarters before it). The data therefore show no reliable step in class E at the quarter a target became known. The drug evidence of a gene usually precedes the first trial of the form and indication that dates the target. In the first snapshot (2016-06-30), 98 of the 136 inflammatory bowel disease targets and 37 of the 145 dated oncology targets already had known-drug evidence. In oncology, 40 targets have a first trial after the first snapshot and no known-drug evidence at that snapshot. Of these, 12 gained known-drug evidence within the archive (6 before their first trial and 6 after it), and 28 never gained it. The oncology lists are for two modalities, while the Open Targets evidence covers the gene in general, which explains most of the 28.

#### Event study: limits

The absence of a shift in class A features does not show that they are free of research-attention bias. The study covers 15 of them over eight quarters. With 39 to 155 events in the target groups, quarterly resolution and one class E feature, the study supports the class A and D assignments. It cannot show the step that defines class E, and it does not allow a choice between dating a target by its first trial and by the registry date. The step of class B in some association groups and not in others is descriptive, and the data do not say why.

#### Neighbourhoods: design

Every gene is a point in the space of the features of a snapshot, each feature taken as the rank percentile among the genes that have a value (a missing value at 0.5), and the ten nearest genes (Euclidean distance) are its neighbours. For each list and each year-end at which at least ten genes were known and at least five were still to become known (90 list–cutoff pairs), we took the share of known genes (P at the cutoff) among the neighbours of the later genes (F) and divided it by the share of known genes among all genes. This enrichment is 1 when the later genes have no more known genes next to them than chance. It was computed with the features of the cutoff and with the features of the latest snapshot (2024-12-31), for five feature sets (classes A, A to B, A to C, A to D and A to E; features that are built from the evidence of the list itself were left out for that list), and also for the known genes themselves and for random genes in no list, in the same number as the later genes. A second measure compares the ten nearest genes of every gene at two snapshots by the Jaccard index of the two sets, over the features that exist in both. The measures describe the data and do not score a method. Intervals are 95% bootstrap intervals over genes (`analysis/neighbour_enrichment.tsv`, `analysis/neighbour_stability.tsv`).

#### Neighbourhoods: observations

At the cutoff, the later genes had a median enrichment of known genes among their neighbours of 2.44 in the space of class A, 3.62 with classes A to C, 5.15 with A to D and 4.86 with A to E. The known genes themselves had 4.49 and 11.2, and random genes 0.83 to 0.88, below 1 because they lie away from the known genes (Fig. 5a). Admitting more classes therefore brings the later genes nearer to the known genes at the cutoff already, with the evidence of that date. Taking the features of the latest snapshot in place of the cutoff changed the enrichment of the later genes little: the median paired change was 0.00 (class A), 0.00 (A to B), 0.17 (A to C), 0.04 (A to D) and 0.25 (A to E), and the enrichment rose in 44% to 67% of the pairs. The change varied widely between lists (*−*32.9 to +44.2), because a list of a few genes moves a lot with one neighbour. One quarter apart (2024-09-30 and 2024-12-31) the neighbours of a gene overlapped by a median Jaccard index of 0.65 in the space of class A and 0.74 with A to E. Four years apart (2016-12-31 and 2020-12-31) the index was 0.09 and 0.05, four years later (2020 and 2024) 0.12 and 0.14, and eight years apart (2016 and 2024) 0.055 and 0.036 (Fig. 5b). The neighbourhoods of the genes thus change with the releases, and more in a space with more classes between 2016 and 2024, so the feature space as of one date is not the space of another.

#### Neighbourhoods: limits

The enrichment depends on the size of a list and on the choice of ten neighbours, and a list with few later genes gives an unstable value. The analysis does not separate the effect of a new release from that of a change in the number of known genes, and it is not a test of leakage into a model.

### 5.6 Coverage across cutoffs

Sources start at different dates, so the features, the genes with data and the known genes that a cutoff offers change with it (Table 7 for the features and Table 8 for the lists; `snapshots/coverage_features.tsv` and `snapshots/coverage_labels.tsv` give every feature set, class and list). The full feature set holds 30 features at the end of 2016 in every area and 66 to 68 at the end of 2024 (66 for inflammatory bowel disease, metabolism and psoriasis, 68 for oncology). The biology-only set grows from 20 to 37 to 44 features. Gene coverage is incomplete before 2023. The gene universe is that of the current Human Protein Atlas, and 18,750 of its 20,151 genes have at least one feature at the end of 2016 and 19,600 at the end of 2020.

The number of held-out genes falls as genes are reached, and it differs by orders of magnitude between lists (Table 8). The immuno-oncology list has two later genes at the end of 2020 and none from the end of 2021, and the cancer driver list has none at the end of 2024. The T-cell-engager list has 3 later genes from 2023. The ADC list has the most later genes among the trial lists of oncology (26 at the end of 2016 and 4 at the end of 2024), and inflammatory bowel disease has 43 and 6. The association lists keep more: the second-study lists of metabolism have 150 to 511 later genes at the end of 2024, the psoriasis list 49, and the ClinGen lists of metabolism none and one. These counts bound what a cutoff can show. A rank statistic over fewer than about ten later genes is illustrative only. Two kinds of interval remain valid at low counts: a bootstrap that resamples the genes, with the gene as the unit, and a permutation of the labels within the candidate genes. `herald_data.leakage.cluster_bootstrap` is an example of the first.

### 5.7 Label dating

Manual audits of the automatically dated trial records ship with the dataset (`curation/, 97` records in two tables). They were checked against the primary ClinicalTrials.gov and drug records. The table `public_label_audit.tsv` (54 records) accepts, rejects or re-dates oncology targets that a single trial supports. The table `target_audit.tsv` (43 records) rejects inflammatory bowel disease drug records whose trial used the drug as co-medication or conditioning and not as a treatment of the disease. It also adds four drug records that the automatic rule missed. These audits target specific edge cases and do not estimate the precision of the automatic matching.

For that purpose, a random sample of the automatically accepted targets was assessed against the trial and drug records shown beside each target (`herald-data labels sample`). The sampled population holds the targets that no manual audit decided, including the single-trial hits that no audit reviewed: 50 of 52 oncology targets and 50 of 134 inflammatory bowel disease targets. The Methods (Audit of the automatic labels) describe the automated review and the hand review of the sample. The sample holds 100 targets, 50 of inflammatory bowel disease and 50 of oncology, and the reviewer judged all of them.

The reviewer judged 93 targets correct, 6 incorrect and 1 unclear. The precision of the labels, correct among correct and incorrect, is 0.94 overall (95% Wilson interval 0.87 to 0.97; 93 of 99), 0.92 for inflammatory bowel disease (0.81 to 0.97; 46 of 50) and 0.96 for oncology (0.86 to 0.99; 47 of 49). If the unclear target counts as incorrect, the oncology precision is 0.94 and the lower bound of the overall interval 0.86 (`analysis/label_audit_judged.tsv`, `analysis/label_precision.tsv`). The sample is random and the reviewer judged all of it, so the interval is an estimate of the precision among the 186 targets that no manual audit decided; it does not cover the audited targets. The six incorrect targets are four in inflammatory bowel disease and two in oncology. Of the four, *FGFR2*, *HRH1* and *IMPDH1* have a drug that served as a conditioning medication, and *MS4A1* has a trial of systemic lupus erythematosus. In oncology, *CD4* is a regulatory T-cell therapy and not an ADC or a T-cell engager, and for *CD7* the drug is an ADC but the trial treats graft-versus-host disease and not cancer. Five of the six are mismatches of the indication and one of the drug modality. The unclear target is *EGFR*: its record names three drugs, two ADCs and one antibody– photosensitizer conjugate, which is not an ADC.

The reviewer and the automated review agreed on 96 of the 100 targets (96%). Of the 98 targets that both judged correct or incorrect, 90 were true positives, none was a false positive, 2 were false negatives and 6 were true negatives. Accuracy was 98.0%, precision 100% (90 of 90; Wilson interval 0.96 to 1.00), recall 97.8%, specificity 100% (6 of 6), the F1 score 0.989 and Cohen’s kappa 0.85.

The areas differ. All 50 inflammatory bowel disease targets agreed. In oncology 46 of 50 agreed, and the accuracy was 95.8% among the 48 decisive targets; 2 more were left out as unclear. The automated review called 8 targets incorrect, 4 in each area. The reviewer confirmed 6 and reversed 2, so a negative verdict was right in 6 of 8 cases (negative predictive value 75%).

All four disagreements were in oncology. The sheet carries no molecule name for 18 of the 50 oncology targets, among them three of these four disagreements and also *CD4* and *CD7*. For *LY75* and *CDH17* the review found no ADC or T-cell engager in the trial, and the reviewer found that the trial named an ADC, for *LY75* one that targets the gene, which the review had missed. For *ADAM9* the review was uncertain about the target relationship, with low confidence, and the reviewer judged it correct because the ADC of the trial targets *ADAM9*. For *EGFR* the review judged the record correct, and the reviewer unclear because one of its three drugs is not an ADC. The modality step failed in three of these four (*LY75*, *CDH17* and the mixed record of *EGFR*), and the relationship step in two (*LY75*, *ADAM9*). Three of the four disagreements carried a high confidence of the automated review, so a high confidence does not mark the records that need a review. The review agreed with the reviewer on 63 of 66 targets at high confidence, 32 of 32 at medium confidence and 1 of 2 at low confidence. The reviewer wrote a note for 45 targets. In 28 the note confirms the evidence sources and conclusions of the review, and in two of these it adds that the review found a ChEMBL record that the reviewer had missed. Three notes record that the review had missed information that the reviewer found (*LY75*, *ADAM9* and *CDH17*).

The reviewer did not see the verdict of the automated review. A single reviewer judged all targets, so the agreement between two reviewers is not known, and an error of the reviewer is an error of the reference. The files `analysis/label_audit_agent_review.tsv` and `analysis/label_audit_agreement.tsv` hold the verdicts, the answer and confidence of each step, and the notes of every target, and the agreement tables by area and by confidence. The evidence that the automated review quoted from ChEMBL, DrugBank and PubMed, and the molecule names of ChEMBL, are not shipped, because the licence of the dataset cannot carry them. The hand-reviewed table of immuno-oncology drugs (`io_targets.tsv`, 76 genes) is shipped beside the audits.

No list of the deposit requires a minimum number of trials. Single-trial targets stay in the lists when an audit confirmed them, and 10 that no audit reviewed stay as well. A stricter rule would remove mostly recent targets, which have had less time to accumulate trials. A rule of at least two trials, or a clinical-stage drug record, would shrink the ADC list from 65 to 55 targets and the T-cell-engager list from 31 to 17 (46 and 11 with three trials). Of the 10 ADC targets lost under the two-trial rule, 8 have a first trial after 2020-02-03, and of the 14 T-cell-engager targets lost, 10 do. At the end of 2020 the ADC list has 13 later targets under the deposited rule, 6 under the two-trial rule and 5 under the three-trial rule. The T-cell-engager list has 10, 0 and 0 (`analysis/threshold_sensitivity.tsv`). A trial-count rule removes later targets as a result of how it is defined, so the deposited lists do not use one.

The registry date and the first-trial date differ for most oncology targets. For 33 of the 64 ADC targets that have a registry date, the registry showed a qualifying trial later than the trial’s start. The median delay was 70 days, 13 targets had more than a year, and the longest was 4,412 days. The same holds for 18 of 31 T-cell-engager targets (median 30 days; 3 more than a year, longest 959 days) and for 43 of 76 immuno-oncology targets (median 75 days; 17 more than a year, longest 3,139 days). For the other targets the two dates are equal or the registry date is earlier, and they count from their first trial. At year-end cutoffs, dating by the registry changes the side of at most two targets of any list. The change is one or two targets at each cutoff for the ADC list, none to two for the T-cell-engager list and none for the immuno-oncology list. The default therefore matters for individual targets and little for list-level counts. None of the other 16 lists has a registry date. Each is dated by its own record (the first trial for the inflammatory bowel disease lists, an approval, a study, a release or a classification), so `known_from` equals the date of the list (`trial_start_only in known_from_basis`). A replay with these lists establishes when the evidence existed and not when a registry or a catalogue showed it.

#### Dates of the GWAS lists

The ten GWAS lists (five diseases, each with a first-study and a second-study list) take their dates from the GWAS Catalog. We compared them with PubMed, an independent record of the same publications (`herald-data citations --check`; `analysis/gwas date check.tsv`). The lists name 434 papers (723 list– paper pairs, 14,778 gene rows). For each pair, the earliest date that PubMed has for the paper (electronic publication or publication) was compared with the date of the list. In 713 pairs PubMed shows the paper on or before the list date, with a median of 113 days between the two (90th percentile 694 days), which is the delay of the Catalog entry. In 10 pairs (8 papers, 35 gene rows, 0.2%) PubMed dates the paper 43 to 167 days after the list date, so the Catalog counts these genes as known before the paper appears in PubMed. We did not establish why. A preprint is one possible reason.

### 5.8 Surface features of the ADC list

An antibody–drug conjugate acts on an antigen at the cell surface, so the genes of the ADC list should differ from other genes in surface features, and the extraction should show it. The list builder requires an extracellular region in UniProt of every candidate (Methods), so a known ADC gene that has a topology value is a candidate by construction, and a comparison with all genes would repeat that rule. The comparisons that carry information are those among the 2,951 candidates of the extraction of 2020-02-03 (genes with an extracellular length above zero), 64 of which are known ADC genes (Fig. 6; `analysis/adc surface check.tsv`). The 65th known gene, *LY75*, has no topology value in that extraction and is left out. Among the candidates, the known ADC genes have a longer largest extracellular segment (median 362 against 117 amino acids; area under the curve 0.72, 95% bootstrap interval 0.68 to 0.76) and fewer transmembrane helices (median 1 against 3; 0.27, 0.24 to 0.31, that is 0.73 for fewer). The UniProt cell-membrane flag does not separate them: 70% of the known genes and 72% of the other candidates carry it (0.49, 0.44 to 0.55). The 19 known genes without the flag all have at least one transmembrane helix. The Human Protein Atlas, an independent antibody-based source of localisation, reports plasma-membrane location for 51% of the 43 known genes that it covers and for 33% of the other candidates (0.59, 0.51 to 0.67). Genes without an extracellular region carry the cell-membrane flag in 8% and the plasma-membrane location in 8%, which shows the effect of the rule itself. The known ADC genes therefore occupy a part of the candidates that has large extracellular regions and mostly few transmembrane helices, as expected for targets of an antibody, and the labels agree with the biology of the modality. The check concerns the labels and the features. It does not show that a method can predict new ADC targets, and the candidate set of the check (the extraction of 2020-02-03) differs slightly from that of the list build, which uses the current topology.

**Figure 6:**
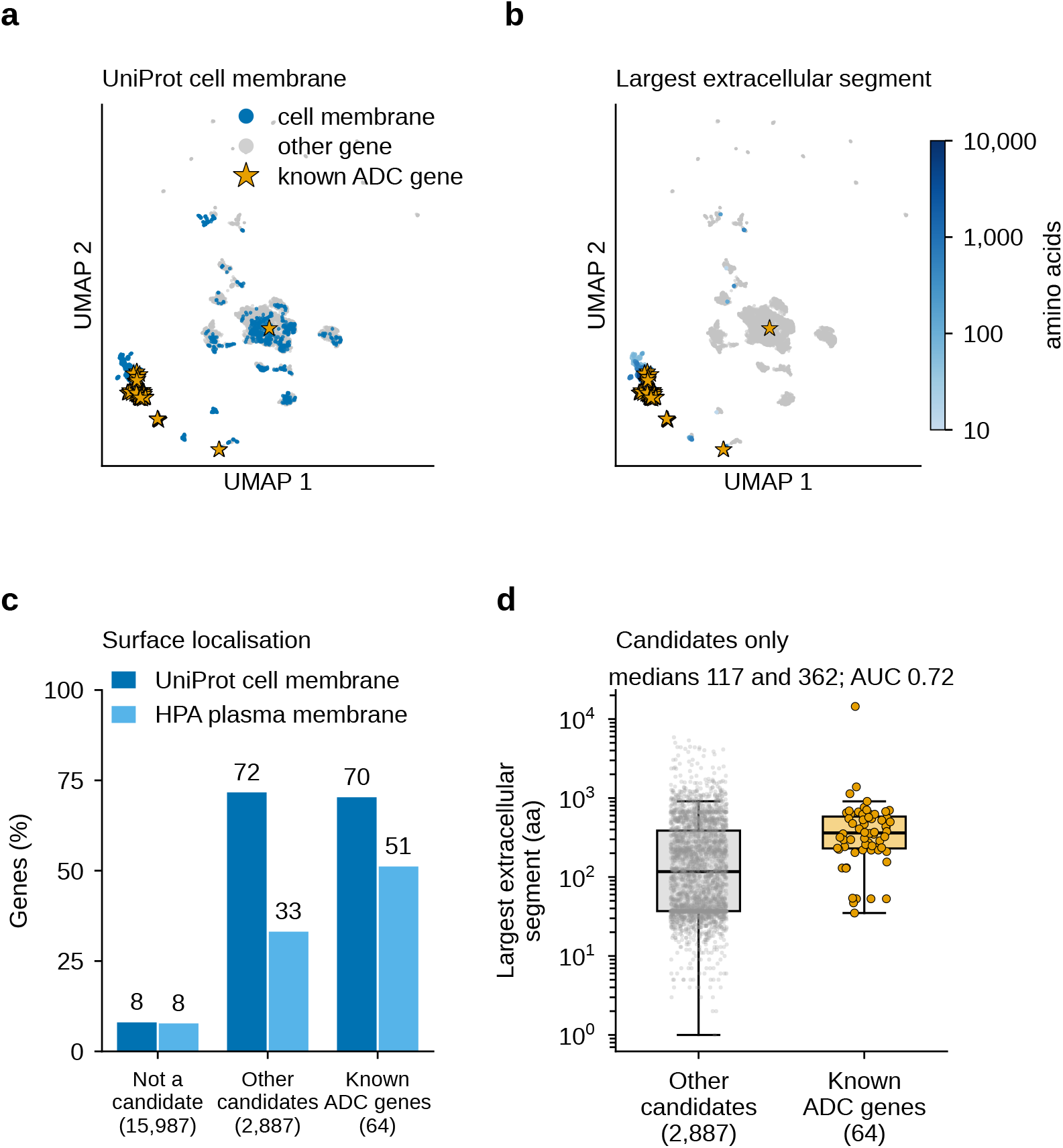
Surface features of the known genes of the ADC list of oncology on 2020-02-03. **a**, **b**, UMAP of the 20,151 genes as in Fig. 2a, with the genes coloured by the UniProt cell-membrane flag (**a**) and by the log of the largest extracellular segment (**b**; grey, no extracellular region or no value). The 65 known ADC genes (orange stars) are drawn on top. **c**, Share of genes with the UniProt cell-membrane flag and with plasma-membrane location in the Human Protein Atlas, for genes that are not candidates of the ADC list (no extracellular region), for the other candidates and for the known ADC genes (number of genes in brackets; the Human Protein Atlas covers fewer genes). **d**, Largest extracellular segment of the candidates (boxes show the quartiles and the median, whiskers the range without outliers; grey points are the 2,887 other candidates and orange points the 64 known ADC genes with a segment value). The text above **d** gives the medians and the area under the curve of the known genes against the other candidates. The candidates are the genes with an extracellular region, which the list requires, so the known genes lie among them by construction.

### 5.9 Identifier mapping

The per-area crosswalk tables map every gene symbol to UniProt, Ensembl and Entrez accessions at build time. For the inflammatory bowel disease area (20,151 genes), 95.8% of symbols carry a UniProt accession, 100% an Ensembl accession and 88.5% an Entrez accession. The oncology, metabolism and psoriasis areas have the same 20,151 genes (the protein-coding genes of the Human Protein Atlas), and the fractions are 95.8%, 100% and 90.5% in each. Symbols without an accession keep an empty field and are not dropped, so every area table shares the same gene universe. `herald-data validate` checks this for the label and feature tables.

### 5.10 Rebuild reproducibility

#### Digests

The manifest records a SHA-256 digest of every deposited file and a tree digest of the files of every archived upstream release. The tree digests cover 183 releases of 20 sources (152,967 files, 133 GB). Regenerating the manifest (`herald-data provenance`) and comparing its digests with the deposited ones verifies a rebuild, except for the structure features described below. The command `herald-data checksums --verify` compares a re-downloaded archive with the recorded tree digests and names the releases that differ.

#### Inputs placed by hand

Three kinds of input cannot be downloaded by a script. They are the DepMap portal releases newer than the last one on figshare (25Q2, 25Q3, 26Q1 and 26Q3), the supplementary tables of three papers (Zecha 2018, Li 2021 and Ito 2020), and the AACT copies of the registry.

#### Withdrawn releases

If a provider withdraws a release, the deposited extractions still serve the cutoffs they cover. A rebuild for another date then uses the nearest earlier release that is still available. This is a fallback reconstruction and not a reproduction of the original snapshot. The record of the extraction names the release used, and a changed file is detected by its digest.

#### Non-deterministic features

Two pocket-volume features of the structure-derived table are estimated by fpocket [42] with Monte Carlo sampling, so they and the digest of that table change on every rebuild. The number of Monte Carlo iterations is the only control, because fpocket offers no random seed on its command line. Between two independent builds, the two features differed for 19,072 of 19,143 genes (median relative difference 1.4%, 95th percentile 4.1%). Every other column of that table, and every other table that the two builds shared, held the same values. Row order and the last floating-point digit of one other structure table can differ between environments. Structure tables are verified by value and not by digest. A user who needs exact numerical reproducibility of the two columns should recompute them.

#### Repeat of the analyses

The event study (`herald-data leakage --events`) was computed a second time from the same snapshots on the same machine. Its three tables were byte-identical to the deposited ones. The two neighbourhood tables (`herald-data neighbours`) were also computed twice and were byte-identical.

#### Tutorial in a new environment

The tutorial notebook was executed a second time in a new Python 3.11 environment that held only the package and the notebook extras (the working environment ran a newer Python). The run raised no error, and the outputs of all 23 code cells and all 8 figures had the same numbers as in the working environment. The UMAP map of the tutorial starts from a principal-component layout and not from the spectral layout of the library, because the spectral start gave different maps in the two environments and the clusters of the map are then not reproducible.

#### Validation checks

`herald-data validate` runs the re-runnable checks. For each disease area it checks that every table the package reads is present. These are the area definition, lists of known genes, crosswalk, labels, negatives, features, roles, gene sets, graph components, STRING and structural tables, eligibility tables and the GWAS annotation table. It checks that every Parquet table opens, that the label and feature tables cover the gene universe of the area, and that no known gene is labelled negative. For the extractions at fixed cutoffs it checks the gene universe, a registry row for every feature, and the absence of features from a source whose licence the release cannot carry. It also checks that the release archive and its calendar are present, that the supplementary calendar entries are complete, and that no non-public file is in the public folder. All checks pass on the current build.

## 6 Usage Notes

Every deposited table is a Parquet, TSV or JSON file and opens with a standard dataframe reader. The manifest allows the integrity of each file to be checked before use. A tutorial notebook ships with the code. It replays the cutoff 2020-02-03 for the ADC list, lists the features available on that day and the genes labelled P and F, and draws a map of the candidate space (as in Fig. 2a).

The time axis and the leakage axis are independent choices, made with `herald-data extract` (Code Availability). The command writes `features.parquet, labels.parquet, feature_registry.tsv` and `extract.json`. The feature set caps the two risk grades (Table 4), and the cutoff selects the releases (Dating of evidence). The biology-only set has fewer features than the others. Both axes should be applied. A biology-only feature set does not replace a cutoff, because the annotation coverage of class A features also grows with the study of a gene (Technical Validation). On the ADC and T-cell-engager lists, evaluation should use genes with an extracellular region, because the flag alone separates their targets from other genes. Dating by the trial registry is the default for the oncology lists. The option `--target-dating trial_start` dates them by their first trial alone. It suits comparisons with analyses that use trial start dates. The two settings differ for at most a few targets at a year-end cutoff (Technical Validation).

### Choosing a list

The question that a method is asked decides the list (Table 8). A ranking of the targets of a kind of drug uses the oncology lists, which are chosen by modality. A ranking of the targets of drugs in a disease uses the trial lists of the inflammatory bowel disease area. A ranking of the genes of a disease uses the GWAS lists and the ClinGen lists of metabolism, and the cancer driver list serves questions about genes of cancer. The features that are made from the evidence of a list give the answer away and should be left out, and `knowledge/evidence_kinds.json` names them for each kind of list (`same_source_features`). For the trial lists no feature is named, and the features of class E, which record drug development, are removed by the no-history feature set.

### Scope of the dataset

The dataset holds public evidence only, so that it can be shared and every analysis repeated. It does not hold the data that a company generates for its own programmes, and a result obtained on it does not describe how a method would perform on such data. Three limits follow. First, the amount of public evidence for a gene follows how much the gene has been studied. The risk grades record this and do not remove it. Second, public resources record what has been published or deposited. Results of programmes that were stopped without publication are mostly absent, and the lists of known genes hold only genes that reached a public milestone. Third, the dates are those of public releases and records. A gene may have been known in a laboratory or a company before the date from which it counts here. The tables are meant for comparing methods under the same dated inputs. They are not meant for predicting how a method will perform on data of another kind.

A cutoff needs the dated release archives. The deposit therefore also holds the extractions at the quarter ends from 2016-06-30 to 2024-12-31 and at 2020-02-03 (`snapshots/`), with the tables of Technical Validation that show which cutoffs support an evaluation. Any other cutoff needs the archive, which one environment variable locates (HERALD_ARCHIVE). The first extraction at a new combination of releases rebuilds the features from those releases and takes correspondingly longer.

The table `literature/gene citations.tsv` supports methods that read documents. For a gene of a list it gives the papers and trials that support it and the date from which each was public. A retrieval system can therefore be given only the documents dated on or before a cutoff and be judged by whether it finds those that made the gene known. The table is not a feature of any ranking. The GWAS and trial lists have citations, and the lists of ClinGen and IntOGen do not (Methods, Citations of the known genes).

The package reads the dataset through one root directory, and one environment variable (`HERALD_DATA _ROOT`) points it at the deposited tree. The disease area is chosen in the same way (`HERALD_AREA`). A new area with the same logic needs an area definition file and its own lists of known genes, and no change to the code.

`herald-data` fetch downloads every upstream release archive (resumable; a fetch that finds its outputs present skips them), and `herald-data` build regenerates the build tables from the releases. A few inputs sit behind a login or a bot check that refuses scripts, and they are placed by hand before the build. For a cutoff, the resolver selects the latest release on or before it from the release calendar, and a cutoff before the first release of a source omits that source. The licence terms of each source apply to the rebuilt tables. The package downloads every source by default, also those that the deposit leaves out, so that a user can rebuild the complete data folder: the ChEMBL release (CC BY-SA 3.0; the list builders read the molecules, mechanisms and targets of the drugs of a modality from it), the optional knowledge graphs (DRKG, Hetionet, PrimeKG), TISCH2 and the sources of the lists (the ClinVar archive, ClinGen, IntOGen, Drugs@FDA and the NCBI gene tables). What a user builds with them is under their licences and is not part of the deposit. The single-cell tables of ten cancers (CHOL, ESCA, HNSC, LAML, LGG, LIHC, OV, PAAD, PRAD and UVM) are not part of the deposit, because TISCH2 [43], which supplies them, states no licence. The deposited tables are those of `herald-data fetch --no-tisch`. A build that includes TISCH2 differs in the oncology area.

## 7 Data Availability

The deposit will be released on Zenodo under the DOI https://doi.org/10.5281/zenodo.23966798 [40] upon publication. The deposit is released under CC BY 4.0 and the `herald-data` code under the MIT licence. The deposit contains the manifest with the release calendar of the source archive, the per-area label, negative, feature, gene-set, graph, crosswalk, eligibility and protein-interaction tables, the structural tables, the lists of known genes, the curation audits and the curated literature selections, the dated citations behind the known genes, the extractions at fixed cutoffs with their coverage tables (`snapshots/`), the feature dictionary and classification rules (`documentation/`) and the tables behind the validation analyses (`analysis/`). The upstream release archives are not part of the deposit. The code below downloads and rebuilds them, and the licence terms of each source apply to them.

## 8 Code Availability

The `herald-data` package is available at https://github.com/huangch/herald-data, release v0.5.0 [44]. The package (Python *≥* 3.10; pandas, NumPy, SciPy, scikit-learn, matplotlib and PyArrow) is distributed under the MIT licence. The commands below download the upstream release archives (a few inputs are placed by hand), regenerate the build tables, write the provenance manifest, run the validation checks, make the folder to deposit and extract a feature set at a cutoff:

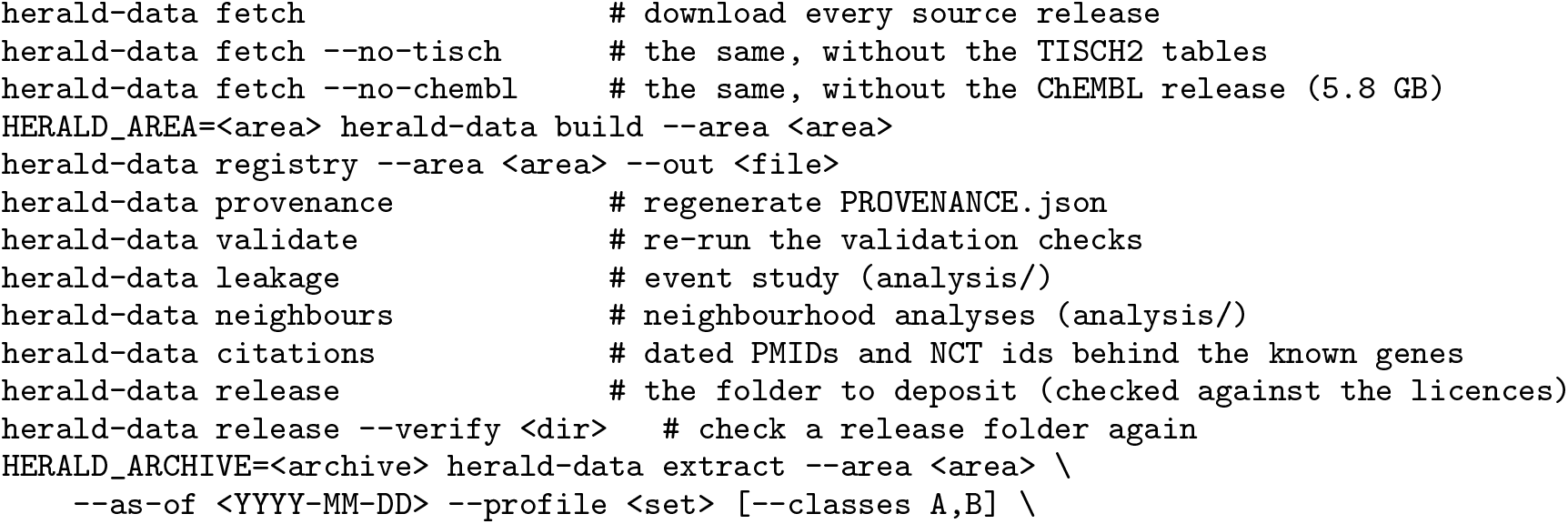
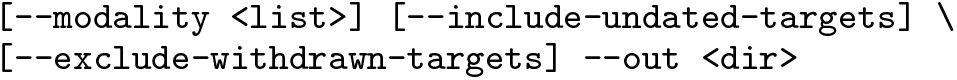

Rebuilding the tables downloads the upstream releases, and the licence of each source applies to them.

